# Substrate availability and dietary fibre regulate metabolism of tryptophan by human gut microbes

**DOI:** 10.1101/2023.06.05.543658

**Authors:** Anurag K. Sinha, Martin F. Laursen, Julius E. Brinck, Morten L. Rybtke, Mikael Pedersen, Henrik M. Roager, Tine R. Licht

## Abstract

Tryptophan is catabolized by gut microbes, resulting in a wide range of metabolites implicated in both beneficial and adverse host effects. However, it remains elusive how the gut microbial tryptophan metabolism is governed either towards indole, associated with adverse effects in chronic kidney disease, or towards indolelactic acid (ILA) and indolepropionic acid (IPA), associated with protective effects in type 2 diabetes and inflammatory bowel disease. Here, we used human fecal cultures in combination with a controlled three-species model to test competition for tryptophan, and measured the resulting metabolites both *in vitro* and in gnotobiotic mice colonized with the three species. We revealed that the generation of specific tryptophan-derived metabolites was not predominantly determined by the abundance of tryptophan metabolizing bacteria, but rather by substrate-dependent regulation of specific metabolic pathways. *In vitro* and *in vivo*, indole-producing *Escherichia coli* and ILA- and IPA-producing *Clostridium sporogenes* competed for tryptophan. Importantly, the fibre degrading *Bacteroides thetaiotaomicron* affected this competition by cross-feeding monosaccharides to *E. coli*, which inhibited indole production through catabolite repression, and thereby made more tryptophan available to *C. sporogenes*, increasing ILA and IPA production. We thus present the first mechanistic explanation for why consumption of fermentable fibres suppress indole production but promote the generation of other tryptophan metabolites associated with health benefits. We conclude that the availability of tryptophan and dietary fibre regulates gut microbiome tryptophan metabolism pathways, and consequently influences the balance between the different tryptophan catabolites generated. This balance has implications for host-microbial cross-talk affecting human health.

## Introduction

Tryptophan is an essential amino acid that is metabolized in the gastrointestinal tract by both host and gut microbiota, resulting in a variety of metabolites, which can affect host metabolism and homeostasis^1,2^. Tryptophan is readily utilized by several gut microbial species, which catabolize it to metabolites including indole, indolelactic acid (ILA), indoleacrylic acid (IAcrA), indolepropionic acid (IPA), indoleacetic acid (IAA), indolealdehyde (IAld), tryptamine etc^1,3,4^. These metabolites regulate host biological processes such as maintenance of epithelial barrier integrity, immune response, protection against pathogens, inflammation and metabolic disorders^1,3,4,5^. Many of them elicit beneficial effects, while others may lead to adverse responses in the host^1^. ILA, IAA and IAld have been shown to stimulate human CD4+T cells to produce IL-22 and reprogramming of intraepithelial CD4+ T helper cells^1,3,6^ thereby promoting tolerance against dietary antigens^7^. IPA also regulates mucosal integrity through the Toll-like receptor (TLR) signaling pathway^8,9^, and is negatively correlated with type 2 diabetes^10,11^, regulates gut permeability^12^, inhibits atherosclerosis^13^ and has antioxidant, anti-inflammatory and neuroprotective properties^14,15,16^. In contrast, indole produced in the gut is converted into the uremic toxin indoxyl sulfate (IS) in the liver. This toxin accumulates in chronic kidney disease (CKD) patients and contributes to the pathophysiology of the disease^1,17,18,19^. Additionally, high indole concentrations in the colon are reported to promote persistent infection with *Clostridium difficile*^20^.

Indole is the most abundant tryptophan metabolite detected in mouse cecal contents as well as in human feces, contributing to 50-75% of the total tryptophan metabolites, and reaching concentrations up to 2.6 mM ^21,22^. Intestinal indole is mainly produced by *Escherichia coli* (*E. coli*) and *Bacteroides* species through a single enzymatic process catalyzed by the *tnaA*-encoded tryptophanase enzyme, which hydrolyses tryptophan into indole, pyruvate, and ammonia^23,24^. Another metabolic pathway, Stickland fermentation, was first described in *Clostridium sporogenes* (*C. sporogenes*), and converts tryptophan into the oxidative pathway product IAA, and the reductive pathway products ILA, IAcrA and IPA (Fig. 1)^25,26,12^. Stickland fermentation is thus a coupled chemical reaction where one amino acid gets oxidized, while another amino acid gets reduced^25^. *C. sporogenes* obtain their energy primarily through Stickland fermentation of amino acids where oxidative metabolism of one amino acid generates ATP via substrate-level phosphorylation, while the redox balance is maintained by reducing another amino acid^25,26,12,27^. Additionally, many *Bifidobacterium* and Lactobacillaceae species are reported to produce ILA from tryptophan in the gut, catalyzed by a specific aromatic lactate dehydrogenase (Aldh) enzyme^3,7,28,29^.

**Fig. 1.**
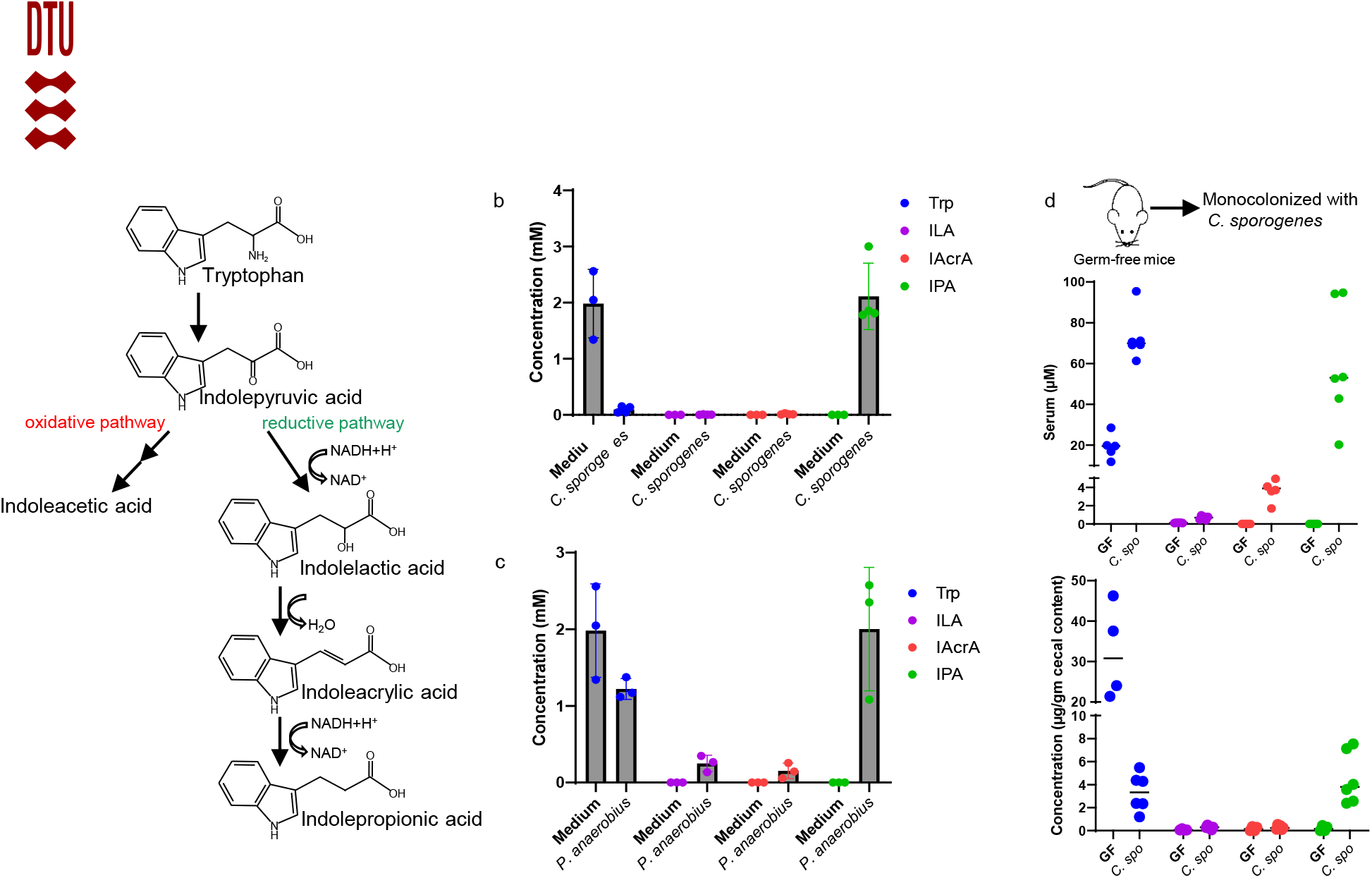
A schematic representation of Stickland fermentation products of tryptophan. Stickland fermentation of tryptophan generates either the oxidative pathway product indoleacetic acid or ILA, IAcrA and IPA through the reductive pathway fermentation^12^.

Despite their important roles in host homeostasis, the factors regulating the generation of these metabolites in the gut remain unknown. The diverse range of tryptophan metabolites produced by the intestinal multispecies community^1^ suggests that bacterial competition for available tryptophan may drive their accumulation in the gut.

Furthermore, recent studies suggest that consumption of fermentable fibre somehow affects microbial tryptophan metabolism. One epidemiological study of five very diverse cohorts reveal that higher daily fibre intake was strongly associated with higher serum level of IPA and lower IS^10^. A positive correlation between serum concentrations of IPA and daily dietary fibre intake is identified in studies of a Finnish population ^11,30^ and a UK cohort^31^. A meta-analysis concludes that serum IS correlates negatively with dietary fibre intake in individuals with CKD^32^ and an intervention with dietary fiber significantly reduce serum IS in haemodialysis patients^33^.

Here, we therefore set out to unravel how production of specific tryptophan metabolites by the intestinal community is affected by the presence of fermentable carbohydrates in the gut environment. We show that tryptophan concentration plays a major role in the accumulation of Stickland fermentation products *in vitro* as well as in human fecal communities. Furthermore, we find that tryptophan availability, degradation of fermentable carbohydrates, and the presence of specific bacterial species govern the balance between tryptophan metabolites formed by Stickland versus tryptophanase pathways *in vitro* and *in vivo*. Our results provide the key mechanisms explaining the observations from multiple human studies.

## Results

### Substrate availability determines tryptophan metabolites production *in vitro* and in human fecal communities

To study specific gut microbial tryptophan catabolism, we used two model species known to perform Stickland fermentation^25,12^. We confirmed *in vitro* that *C. sporogenes* and *Peptostreptococcus anaerobius* (*P. anaerobius*) produced the specific tryptophan metabolites, ILA, IAcrA and IPA, resulting from the reductive pathway of the Stickland fermentation, while tryptophan was consumed (Fig. 1 and Supplementary figure 1a-b). In contrast to germ-free (GF) mice, mice mono-colonized with *C. sporogenes* contained the tryptophan metabolites in cecum and serum (Supplementary Fig. 1c), confirming the strict microbial origin of the metabolites^34^. In addition, cecal concentrations of tryptophan were significantly reduced in *C. sporogenes* colonized mice as compared to GF mice.

Further, since tryptophan was completely consumed by *C. sporogenes in vitro* (Supplementary figure 1a), we hypothesized that levels of Stickland fermentation products depended upon the availability of this substrate. Indeed, we observed clear dose dependent higher accumulation of ILA and IPA in the culture supernatant of both *C. sporogenes* and *P. anaerobius* upon tryptophan supplementation in the medium (Fig. 2a-b) implying that tryptophan availability drives the production of ILA and IPA. Next, we assessed whether higher carbohydrate availability affected production of ILA and IPA, considering that *C. sporogenes* does ferment carbohydrates, although they are not essential for growth of this species if amino acids are present in the environment^25^. No significant change in the tryptophan metabolites were observed upon supplementation of 5 to 10-fold higher glucose in the growth medium (Supplementary Fig. 2a-b), suggesting that Stickland fermentation is unaffected by presence of carbohydrates in the environment.

**Fig. 2.**
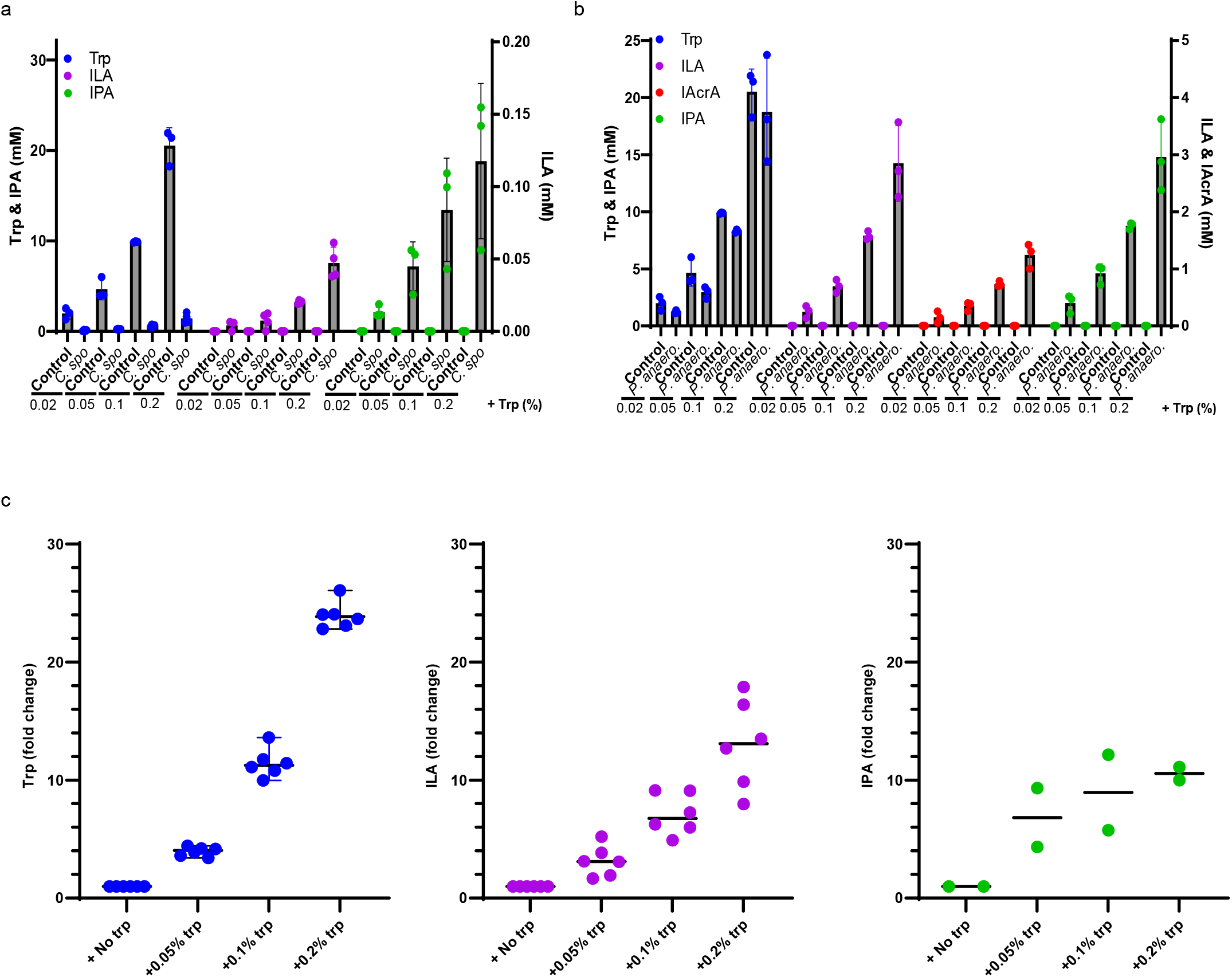
Tryptophan supplementation increases tryptophan-derived Stickland fermentation products. (a) Tryptophan, ILA and IPA accumulation in the culture supernatant of *C. sporogenes* grown in mGAM medium supplemented with final concentrations of 0.02, 0.05, 0.1 or 0.2 % free tryptophan. The left Y axis represents values for tryptophan (Trp) and for indolepropionic acid (IPA) whereas the right Y axis represents indolelactic acid (ILA). (b) Tryptophan metabolites in the culture supernatant of *P. anaerobius* grown in mGAM medium supplemented with final concentration of 0.02, 0.05, 0.1 or 0.2 % free tryptophan. (c) Normalized tryptophan metabolites in the culture supernatant of faecal microbiotas. Six infant faecal microbiotas were cultured either in YCFA medium or YCFA supplemented with 0.05 or 0.1 or 0.2 % of free tryptophan. Specific metabolite concentrations are normalized to the basal level of the given metabolite in the growth medium without tryptophan supplementation. Absolute values of individual faecal cultures are shown in the Supplementary figure 2.

Next, we investigated the effects of substrate availability on tryptophan metabolite production in a complex microbial community. Six infant fecal samples from a previous study^3^ were selected based on the presence of ILA producing *Bifidobacterium* species^3^ or ILA and IPA producing *P. anaerobius*^12^. In agreement with the mono culture experiments, tryptophan supplementation significantly increased ILA and IPA production in the complex communities (Fig. 2c). 16S rRNA amplicon sequencing revealed that different tryptophan supplementation did not lead to noteworthy differences in the individual community composition, suggesting that the increase in ILA and IPA was driven by higher substrate availability and not attributed to a change in the abundance of producer species (Supplementary Fig. 2d).

### Carbohydrate availability affects microbial tryptophan metabolism in fecal cultures

While the ILA, IAcrA and IPA are generated by stepwise reductive Stickland fermentation performed only by a few specific members of the human gut microbiota, indole is produced from tryptophan in a single catabolic step by all bacteria encoding the tryptophanase enzyme gene *tnaA* (Fig. 3a)^23,24,35^. Intestinal indole is mainly produced by *E. coli*^23^. In this species, the *tnaA* gene is under control of carbon catabolite repression and its expression is thus inhibited by the presence of simple carbohydrates such as glucose, arabinose and pyruvate^35,36^. We therefore hypothesized that addition of simple carbohydrates to the growth medium would inhibit indole production in the fecal culture. To test this, we cultured a fecal sample, confirmed by 16S rRNA amplicon sequencing to contain *Escherichia* species, in YCFA medium with low to high concentrations of glucose, maltose and cellobiose as per protocol^37^ (Fig. 3b). Indeed, at low carbohydrate concentrations, indole was readily produced while at higher concentrations, its production was completely inhibited, which confirmed our hypothesis (Fig. 3b) (Supplementary Fig. 3a). 16S rRNA amplicon confirmed that a high abundance of *Escherichia* was maintained even after higher carbohydrate supplementation (Supplementary Fig. 3b). Thus, we conclude that supplementation of carbohydrates inhibited indole production in the complex community without altering the abundance of the producing species.

**Fig. 3.**
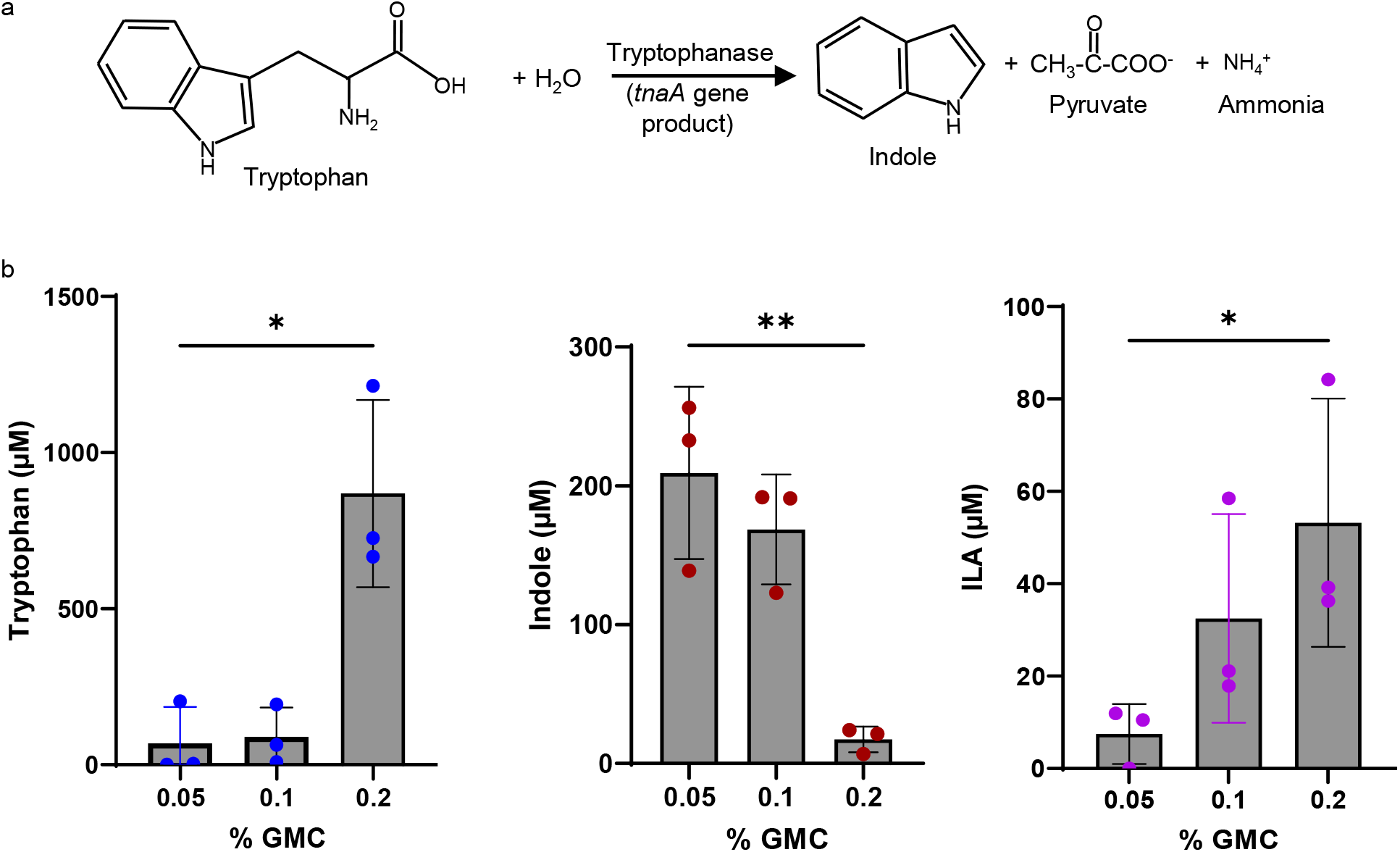
Carbohydrate supplementation inhibits indole production by infant gut microbiota. (a) Schematic representation of tryptophanase-mediated catabolism of tryptophan to produce indole, pyruvate and ammonia. (b) Tryptophan metabolites in the culture supernatant of faecal microbiota in YCFA medium supplemented either with 0.05, 0.1 or 0.2 % glucose (G), maltose (M) and cellobiose (C), collectively referred to as GMC. One infant faecal sample (23.11) was selected for cultivation in three replicates since it contained *P. anaerobius*, capable of Stickland fermentation, and *E. coli*, an indole producer. Metabolites in the individual culture supernatant were normalized against the final OD_600_ of the culture. Results are mean ± SD of three independent experiments. Statistical analysis was done using a two-tailed unpaired t-test. *P < 0.05, **P < 0.01, ***P < 0.001. Individual replicates and their 16S rRNA profile are shown in Supplementary figure 3.

Importantly, inhibition of indole production concomitantly caused more tryptophan (substrate) to remain available in the supernatants from cultures supplemented with 0.2 % carbohydrates (Fig. 3b). Higher tryptophan availability led to increased production of ILA (Fig. 3b) in line with the monoculture and fecal culture experiments (Fig. 2). Oppositely, supplementation with limited amounts (0.05 %) of carbohydrates resulted in more tryptophan conversion into indole and reduction in ILA production (Fig. 3b). Thus, we observed clear inverse correlation between indole and ILA accumulation. The supplementation experiments in complex communities confirm that microbes compete for available tryptophan to produce either indole or Stickland fermentation products, and that the outcome of this competition is influenced by the availability of carbohydrates in the environment.

### Fibre modulates tryptophan availability by inhibiting indole production through cross-feeding

Because simple sugars from the diet do not reach the colonic microbes, we addressed whether catabolism of complex fibres by gut microbes would cross-feed simple sugars that infer catabolite repression in *E. coli*, and thereby affect indole production. For this, we constructed a simple microbial community comprising the three model species; *E. coli* (indole producer); *Bacteroides thetaiotaomicron* (*B. theta*, indole producer^24^, pectin degrader) and *C. sporogenes* (producer of Stickland fermentation products) (Fig. 4a). Measurements of the mono-culture supernatants from *E. coli*, *B. theta* and *C. sporogenes* revealed that *B. theta* indole production was almost 4 to 5-fold lower than that of *E. coli* (Supplementary Fig. 4a). When all three species were co-cultured, only presence of *E. coli* resulted in indole accumulation in the culture supernatant (Supplementary Fig. 4b), revealing *E. coli* as the main indole producer in the defined community. The defined three-species community was cultured in low (0.02%) tryptophan and high (0.05%) tryptophan containing media. The media were further either supplemented or not supplemented with apple pectin. In agreement with observations from mono-cultures (Fig. 2), a 2.5-fold higher supplementation of tryptophan resulted in 2 to 3-fold higher levels of ILA and IPA in the three-species community, confirming that substrate availability determines the Stickland fermentation (Fig. 4b). Furthermore, presence of pectin in the growth media consistently inhibited indole production by 40-50% compared to when pectin was not added, both in the low and in the high tryptophan groups (Fig. 4b). In contrast, tryptophan and ILA both increased in the presence of pectin, and the same trend was observed for IPA (Fig. 4b). Pectin availability thus directed tryptophan metabolism towards less indole production and increased Stickland fermentation in the three species system.

**Fig. 4.**
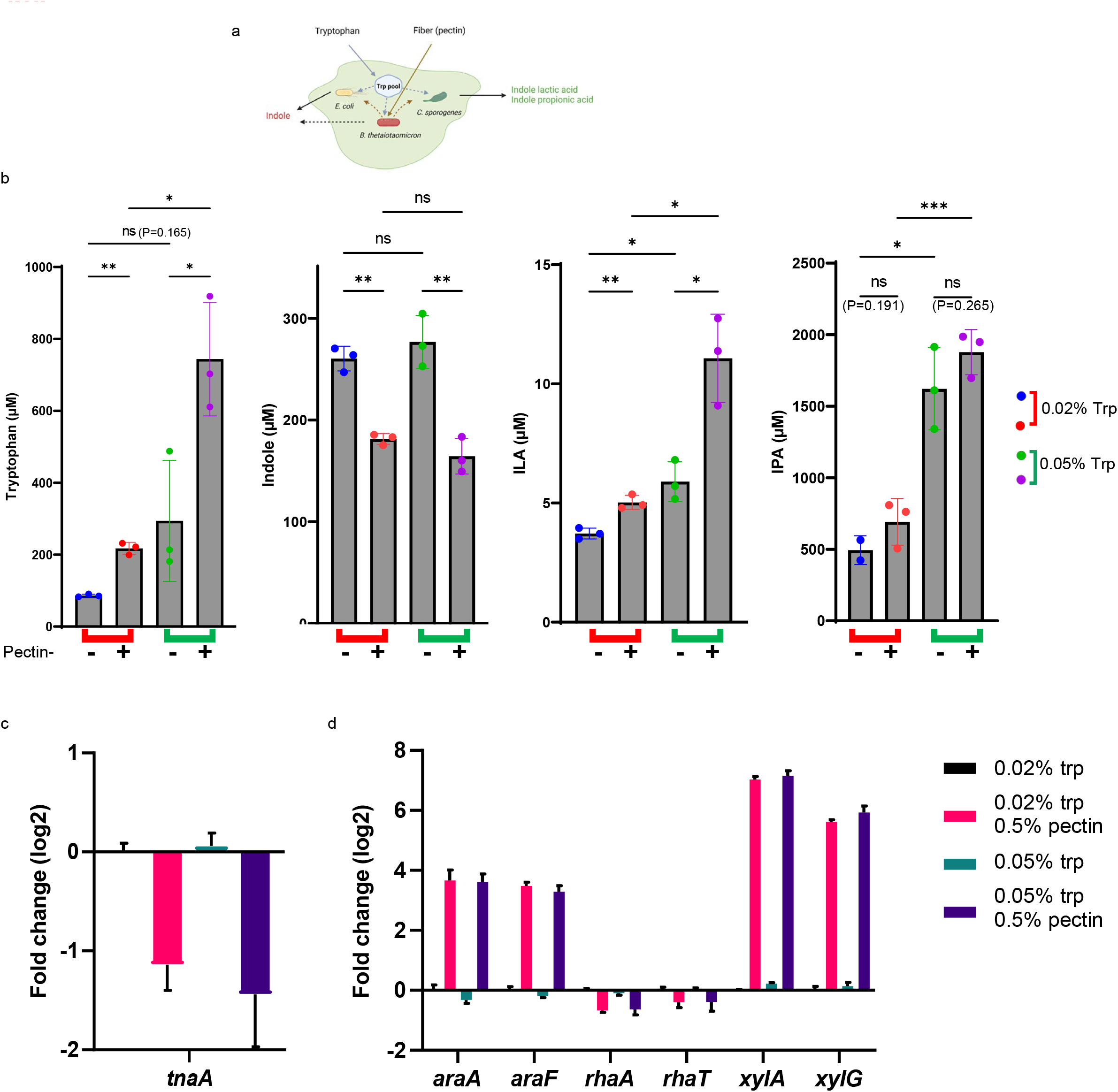
Tryptophan and fibre supplementation alters tryptophan metabolites production. (a) Schematic representation of the bacterial species comprising the defined community. *E. coli* was selected as major indole producer, *B. theta* was selected as fibre degrader and *C. sporogenes* to produce Stickland fermentation products. (b) Tryptophan metabolites in the supernatant of the defined community cultured in mGAM medium supplemented with either 0.02 or 0.05 % free tryptophan. Both low and high tryptophan media were further supplemented with either no apple pectin or 0.5 % apple pectin. (c) RT-qPCR targeting *tnaA* mRNA in *E. coli* in response to pectin supplementation. (d) RT-qPCR targeting mRNAs of arabinose utilizing genes (*araA* and *araF*), rhamnose utilizing genes (*rhaA* and *rhaT*) and xylose utilizing genes (*xylA* and *xylG*) in *E. coli* in response to pectin supplementation. Total RNA was extracted from early stationary phase cultures (∼ 1 OD) and mRNA levels were measured as described in methods. Results are mean ± SD of three independent experiments. Statistical analysis was done using Welch’s ANOVA test in panel b. *P < 0.05, **P < 0.01, ***P < 0.001.

Monosaccharides such as arabinose are known to repress *tnaA* gene expression in *E. coli*^35, 36^. To test whether cross-feeding of monosaccharides resulting from *B. theta* pectin degradation repressed *tnaA* gene expression in *E. coli*, we monitored messenger RNA abundance by reverse transcription quantitative PCR (RT-qPCR) in the defined three-species community. Indeed, when the community was grown in presence of pectin, *E. coli tnaA* gene expression was inhibited 2 to 4 fold (Fig. 4c), explaining the inhibition of indole production and tryptophan consumption. Further inhibition of the *tnaA* gene in both *E. coli* and in *B. theta* was observed when samples were collected after 24 hrs fermentation (Supplementary Fig. 4c). Interestingly, arabinose and xylose utilizing genes were upregulated by 16 to 64 fold, respectively, in *E. coli* in the presence of pectin, suggesting that uptake of these monosaccharides were increased in *E. coli* due to cross-feeding with products of pectin degradation (Fig. 4d). This is in agreement with a previous study showing that *B. theta* digests pectin and upregulates arabinose-, xylose- and rhamnose-utilizing genes in *E. coli*^38^.

These findings demonstrate that pectin degradation results in cross-feeding of simple carbohydrates to *E. coli*, which due to catabolite repression inhibits expression of *tnaA*. Consequently, the conversion of tryptophan into indole is inhibited, making more tryptophan available for *C. sporogenes* for Stickland fermentation.

### Fibres and substrate influence microbiota function to regulate microbial metabolites concentration *in vivo*

Having explored the relations between substrate availability and microbial tryptophan metabolism *in vitro*, we investigated whether substrate availability affected circulating tryptophan metabolites *in vivo*. Four groups of germ-free mice (n=5 per group) were dosed with the three species community and fed for 2 weeks with either normal (2g/kg) or high (16g/kg) tryptophan, in combination with either no pectin or 50g/kg pectin (Figure 5a and Supplementary Fig. 5). There was no significant difference in the total cecal and colonic bacterial load amongst the groups (Supplementary Fig. 5d). However, pectin consumption reduced the relative and total abundance of *C. sporogenes* in both cecum and colon, whereas relative and total abundances of *E. coli* and *B. theta* were similar between the groups (Fig. 5b and Supplementary Fig. 5e).

**Fig. 5.**
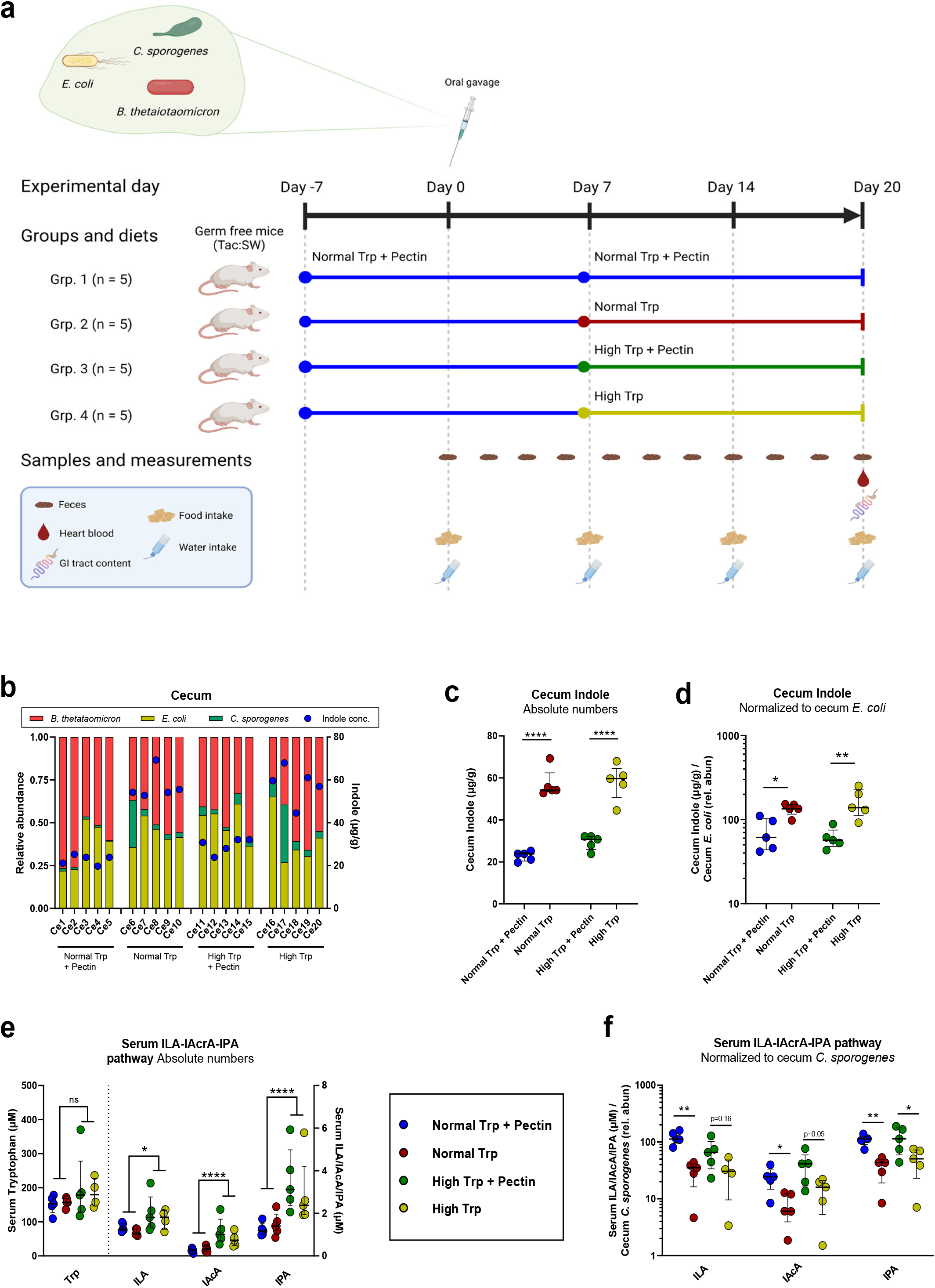
Tryptophan and fibre supplementation modulates tryptophan metabolites production by a defined community in vivo. (a) Schematic representation of experimental plan to evaluate the effect of dietary tryptophan and pectin on tryptophan metabolite production *in vivo*. Germ free mice were placed in four groups (N=5 per group), and fed a diet containing 2 g/kg tryptophan and 50 g/kg pectin for seven days for adaptation. They were then orally gavaged with a mixed culture of *E. coli*, *B. theta* and *C. sporogenes* in equal amounts (OD_600_) and remained for another seven days on the same diet for stabilization. Diets were then changed and mice were fed either a diet with either 2 g/kg or 16 g/kg tryptophan, with or without 50 g/kg pectin for two more weeks. Samples were collected as shown in the scheme. (b) 16S rRNA gene sequencing profiles show the composition of the defined community in caeca of the four groups, overlaid indole values measured in the individual caeca. (c) Absolute caecal indole concentrations. (d) Indole concentration in the caeca, normalized to the relative abundance of *E. coli*. (e) Absolute concentrations of tryptophan, ILA, IAcrA and IPA in serum. (f) Serum tryptophan metabolites (ILA, IAcrA and IPA) normalized to *C. sporogenes* relative abundance in cecum. For plots in panel b-f, lines and error bars indicate median and IQR. Statistical analysis was done across groups within each metabolite measured using One-way ANOVA (panel c) or Kruskal Wallis tests (panel d-f), using uncorrected Fisher’s LSD or Dunn’s posthoc tests to compare between individual groups. *P < 0.05, **P < 0.01, ***P < 0.001. For panel e and f, one value for tryptophan and ILA was excluded as an extreme outlier (Grubbs test, alpha < 0.01).

In agreement with the *in vitro* results, indole concentrations in cecum and colon were consistently lower in the presence of pectin in the diet (Fig. 5b-d and Supplementary Fig. 5e-g). Notably, dynamics of indole accumulation was not related to the relative abundance of *E. coli* in the individual mice (Fig. 5b and Supplementary Fig. 5e). Normalization of indole concentrations to the abundance of the producing *E. coli* in each animal thus confirmed that the indole-reducing effect of dietary pectin was explained by a reduction of the production of indole from each bacterial cell, rather than by a decrease in the number of producing cells (Fig. 5d and Supplementary Fig. 5f-g). Cecum and colon IPA concentrations were very low (Supplementary figure 5h-i), and ILA was below detection level in intestinal samples, suggesting that these compounds are efficiently absorbed from the gut.

In serum, we found higher concentrations of ILA, IAcrA and IPA in mice fed with the high tryptophan diets, confirming that increased tryptophan availability led to increased Stickland fermentation *in vivo* (Fig. 5e). However, there was no significant difference between the absolute serum concentrations of tryptophan among the groups (Fig. 5e), probably due to a tight host regulation of circulating free tryptophan concentration in the serum as reported earlier^22,39^. ILA, IAcrA and IPA concentrations in serum did not follow the abundance of *C. sporogenes* in cecum or colon of individual animals. However, normalization of serum metabolite concentrations to the *C. sporogenes* relative abundance in cecum and colon suggest that each *C. sporogenes* cell produced more ILA, IAcrA and IPA in the presence of pectin (Fig. 5f and Supplementary Fig. 5j), indicating an upregulation of the tryptophan Stickland fermentation pathway in pectin fed mice. A similar picture was not seen for Stickland fermentation of other substrates, such as valine and leucine, since we observed a clear correlation of their Stickland reaction products, isovaleric acid and isobutyric acid, with *C. sporogenes* abundance independent of diets (Supplementary Fig. 5k). This confirms that the increased amount of tryptophan Stickland fermentation metabolites was not explained by a general increase in the producing strains, but by an increased amount of tryptophan available to bacterial fermentation in animals fed with pectin (Fig 6).

**Fig. 6.**
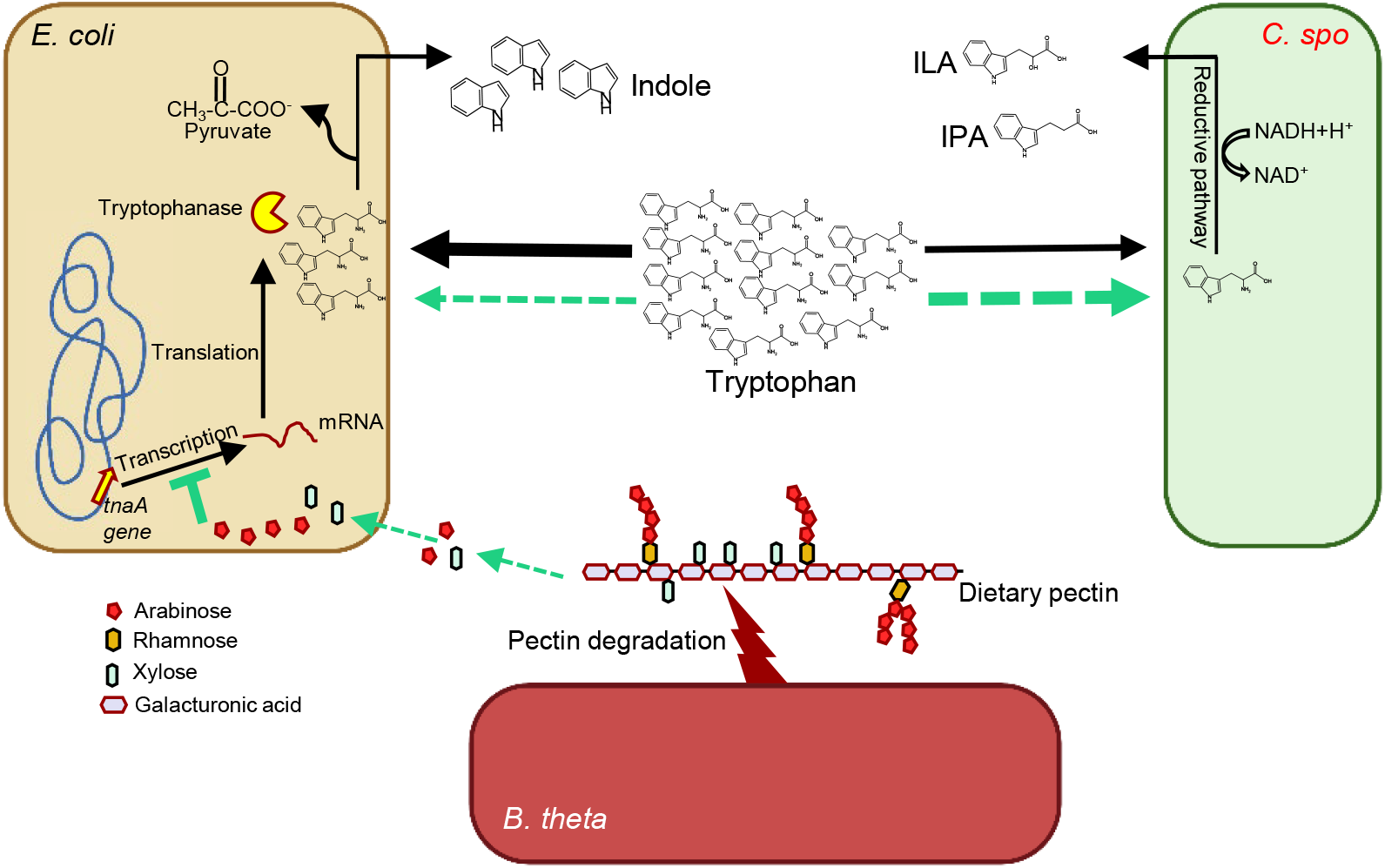
Dietary fibre and substrate mediated impact on tryptophan metabolism. In the gut, multiple bacterial species require tryptophan for their metabolism, and produce bioactive molecules important for host health. *Escherichia coli* catabolises tryptophan into indole to generate pyruvate, while *C. sporogenes* regenerates NAD+ and produces indolelactic acid (ILA) and indolepropionic acid (IPA) through the Stickland fermentation reductive pathway. The fibre degrader *B. thetaiotaomicron* degrades pectin, and thereby release monosaccharides available to *E. coli*. The monosaccharides represses expression of the *E. coli tnaA* gene encoding tryptophanase, thereby making more tryptophan available to Stickland fermenters in the gut environment. Black arrows show events occurring in the absence of fibre, while green arrows designate events preferentially occurring in the presence of fibre. Thick and thin arrows depict enhanced and reduced flow of tryptophan, respectively.

## Discussion

We show that microbial competition for tryptophan determines which tryptophan metabolites are produced by communities of intestinal microbes, and that this competition is significantly affected by the availability of tryptophan as well as of simple carbohydrates originating from fibre degradation. We propose a model explaining how dietary fibres influence microbiota activity and thereby alter the balance between formation of beneficial (ILA, IAcrA, IPA) and potentially adverse (indole, precursor of indoxyl sulphate) tryptophan metabolites (Fig. 6).

In the gut, ILA, IAcA and IPA is generated by Stickland fermentation of tryptophan, performed e.g. by *C. sporogenes* and *P. anaerobius*. Indole, on the other hand, is generated from tryptophan by bacteria expressing the *tnaA*-encoded tryptophanase enzyme, exemplified by *E. coli*.

We reveal that increased levels of substrate leads to a high product formation of the tryptophan catabolizing Stickland fermentation *in vitro* and *in vivo*. Additionally, we show that carbohydrates inhibit the conversion of tryptophan to indole by downregulation of *tnaA* gene expression through catabolite repression. This leads to more available tryptophan for bacteria performing Stickland fermentation. Higher tryptophan availability to the Stickland fermenting gut microbes can thus be achieved by two means: Either by including more tryptophan in the diet, or by inhibiting the consumption of tryptophan by the indole producing species through increasing intestinal carbohydrate levels by consumption of dietary fibre (Fig. 6).

In agreement with our model, indole production has previously been shown to be inhibited in the presence of starch during batch fermentation using human fecal slurries^40^, and pectin and inulin supplementation has been reported to decrease indole and increase ILA, IAA and IPA accumulation in the medium of a batch culture inoculated with human microbiota^41^. Additionally, a study on pigs aiming to understand the effect of feeding with non-starch polysaccharides (NSP) revealed that the intestinal amounts of indole was lower while IPA and IAA tended to be higher in pigs fed a high-NSP diet^42^.

Our model (Fig. 6) offers a key to the mechanistic understanding of results obtained in several previous human studies, reporting that dietary fiber intake correlates positively to beneficial tryptophan metabolites such as ILA and IPA, but negatively to the potential deleterious metabolites such as indoxyl sulphate^10,11,30,42^.

Abundance of *E. coli* increases significantly during chronic kidney disease (CKD), explaining high levels of indole in the gut, which leads to high levels of indoxyl sulfate in the serum of CKD patients^23^. Increased indoxyl sulfate may contribute to not only kidney damage and renal insufficiency, but also to atherosclerotic lesions observed in dialysis patients^43^. Also, the intestinal pathogen *C. difficile* actively upregulates high indole production by indole-producing gut microbes that allow *C. difficile* to proliferate and cause persistent infection^20^. Thus, it is desirable to down-regulate the gut microbial indole production. Limiting protein intake and increasing dietary fibre intake is reported to reduce serum indoxyl sulphate, and has been considered as an treatment option for CKD^43,44,45^.

Our model provides a rational for directing gut microbial tryptophan metabolism away from indole production and towards generation of beneficial Stickland fermentation products including ILA^1,2,3^, IAcrA^1,4^ and IPA^10,13,46,47^ through dietary intervention. It is worth to highlight that unlike most previous approaches, this model builds on alteration of microbial activity and gene regulation rather than alteration of microbiota composition and/or abundance of specific producer species. We believe that the future of microbiome research lies in including microbial metabolic activity, not only by assessing abundance of bacterial genes, but also the cues and triggers that regulate their expression. Supporting this believe, a recent analysis of a Dutch population-based cohort revealed a striking lack of correlation between metagenomics gene abundance and corresponding microbial metabolites^48^. Our finding that interspecies competition for a specific substrate in combination with catabolite repression determines levels of relevant microbial metabolites in the gut can most likely be extrapolated to many other substrates and competition/cross-feeding interactions of the gut microbiome, which remains to be revealed.

## Methods

### Bacterial strains and media

Representative bacterial strains *Clostridium sporogenes* (DSM 795), *Peptostreptococcus anaerobius* (DSM 2949), *Bacteroides thetaiotaomicron* (DSM 2079) and *Escherichia coli* K12-MG1655 (DSM 18039) were purchased from DSMZ (German Collection of Microorganisms and Cell Cultures GmbH, Germany). They were revived on Gifu Anaerobic medium, Modified (mGAM) agar plates, glycerol stocks were prepared and stored in −80°C until further used. For batch culture experiments, bacterial strains were revived on mGAM agar plates and grown overnight as primary cultures in mGAM broth medium under mild shaking condition. Next morning, they were then diluted again as 0.02 OD_600_ into 3 ml mGAM broth medium as secondary cultures and grown for 48-72 hrs in mild shaking condition. Each strain was cultured at least in triplicates. Culture medium without inoculation were used as controls. After 72 hrs of fermentation, OD_600_ was measured and samples were put on ice. 1 ml each of the samples were centrifuged at 14000 rpm for 10 minutes at 4°C, supernatants collected and stored at −20°C. For supplementation experiments, media were prepared with different amount of amino acids as indicated in the figures, autoclaved and used as described above. Fecal microbiota was cultured in YCFA medium supplemented with 0.2 % each of glucose, maltose and cellobiose, as described earlier to support large group of gut bacterial species^37^. Collected supernatant samples were processed for metabolites extraction and analysis as described in the next section below. Pectin from apple for defined community *in vitro* experiments was purchased from Sigma-Aldrich, Merck (93854-100G). All growth experiments were performed inside Whitley A95 anaerobic workstation maintained at 37°C and all the plates or media were incubated inside the workstation at least 24 hrs before use to maintain anoxic conditions.

### Infant fecal samples

Six infant fecal samples were selected from The Copenhagen Infant Gut (CIG) cohort obtained for a separate study in the lab with approval from The Data Protection Agency and from the Ethical commitee^3^.

### Animal experiments

All germ free (GF) Swiss Webster mice (Tac:SW) used for experiments were bred in the GF facility at the National Food Institute, Technical University of Denmark, maintained on an irradiated chow diet (Altromin 1314, Brogaarden ApS, Lynge, Denmark) and transferred to experimental isolators before experiments began. In all experiments the environment was maintained on a 12h light/12h dark cycle at a constant temperature of 22 ± 1 °C, with an air humidity of 55 ± 5% relative humidity and air was changed 50 times/hour. For all experiments, the GF status of mice prior to oral gavage was confirmed by inoculation of feces from all groups separately into BHI broth (25°C and 37°C, aerobic incubation), mGAM broth (37°C, anaerobic incubation) and plating on blood agar (37C, aerobic incubation) and evaluation after 24h and 2 weeks of incubation. All animal experiments were approved by the Danish Animal Experiment inspectorate (License Number: 2020-15-0201-00484) and were overseen by the National Food Institute’s in-house Animal Welfare Committee for animal care and use.

### Mono-colonization experiment

Six GF SW mice (Nmales = 2, Nfemales=4) were at the age of approximately 6 weeks, transferred into an experimental isolator and housed individually (Makrolon Type II cage, Techniplast, Varese, Italy) with bedding, nesting material, a hiding place and a wooden block. Mice had free access to sterile drinking water (Glostrup Hospital, Denmark), and were maintained on a standard purified diet containing 0.21% Tryptophan (D10012G, Research diets, New Brunswick, NJ, US) throughout the experiment. All mice were acclimatized for 7 days, before oral gavage with a 200 µL PBS-washed *C. sporogenes* culture (grown overnight in mGAM medium and washed twice with PBS) and euthnization after four days of colonization. Four female GF SW mice maintained on the same diet for at least 7 days were used as controls for detection of tryptophan catabolites.

*C. sporogenes*-colonized as well as GF control mice were anestisized in hypnorm/midazolam (0.1 ml/10g SC), terminal heart blood (*C. sporogenes* colonized mice) or portal vein blood (control mice) was collected and the mice were euthanized by cervical dislocation, before collection of cecum content. Serum was generated from the blood samples after 30 min of coagulation, centrifugation (2000xg, 10 min 4°C) and aspiration of supernatant into Eppendorf tubes stored at −20°C until further processing. Cecum content was homogenised 1:4 with sterile MilliQ water by vortexing and subjected to centrifugation (10000xg, 5 min 4°C), before collection of both pellet and supernatant in separate tubes snap frozen on dry ice and stored at −80°C until further processing. The primary outcomes assessed in this experiment was detection/quantity of tryptophan and tryptophan catabolites produced by the Stickland fermentation pathway (IAA, ILA, IAcrA and IPA) in cecum content and blood.

### Dietary tryptophan and pectin

Twenty GF SW mice (Nmales = 9, Nfemales=11) were at the age of approximately 10 weeks pseudo randomized intro groups based on gender and transferred into 4 separate experimental isolators (each experimental isolator contained a group of 5 mice, including either 3 males + 2 females or 3 females + 2 females). Mice were housed individually (Makrolon Type II cage, Techniplast, Varese, Italy) with bedding, nesting material, a hiding place and a wooden block. All mice were acclimatized for 7 days before oral gavage with the defined community of bacteria. All mice has free access to sterile drinking water (Glostrup Hospital, Denmark) and were maintained on an irradiated purified diet named “Normal Trp + Pectin” (A22033102-1.5V, Research diets, New Brunswick, NJ, US) consumed ad libitum from day −7 to day 7. On day 7 all, but group 1, shifted to diets containing either “Normal Trp” (A18041301R-1.5V, group 2), “High Trp + Pectin” (A22033103-1.5V, group 3) or “High Trp” (A22033101-1.5V, group 4), whereas group 1 continued on “Normal Trp + Pectin” until the experiment ended on day 20 (see all diet compositions in Supplementary table 1). On day 0 all mice were individually orally gavaged with a 200 µl PBS-washed mixture of individually cultured *B. theta*, *C. sporogenes* and *E. coli*. These species were cultured individually in mGAM medium for overnight, centrifuged and cell pellet were washed twice with PBS. Equal OD cells of all three species were then mixed and prepared for gavage. Further, 16S rRNA amplicon sequencing for DNA obtained from cecal luminal content, confirmed that the mice were colonized only by the three inoculated species. From day 0, fresh fecal samples were obtained every second day and water and food consumption was registered for each individual mouse weekly (day 0, day 7, day 14 and day 20).

At day 20 mice were anestisized in hypnorm/midazolam (0.1 ml/10g SC), terminal heart blood was collected and the mice were euthanized by cervical dislocation, before collection of gastrointestinal luminal content and tissue. Serum was generated from the heart blood after 30 min of coagulation, centrifugation (2000xg, 10 min 4°C) and aspiration of supernatant into Eppendorf tubes stored at −20°C until further processing. Cecum and colon luminal content homogenised 1:4 with sterile MilliQ water by vortexing and subjected to centrifugation (10000xg, 5 min 4°C), before collection of both pellet and supernatant in separate tubes snap frozen on dry ice and stored at −80°C until further processing. The primary outcomes assessed in this experiment was detection/quantity of tryptophan and tryptophan catabolites produced by the Stickland fermentation pathway (IAA, ILA, IAcrA and IPA) as well as indole in cecum and colon content as well as blood.

### Statistics

All statistics for the animal experiments were performed with the GraphPad Prism software (v9.5.0). Normal distributions were evaluated by the Shapiro-Wilk test. Water intake, food intake and estimated tryptophan intake (food intake * trp content of diets) were compared between groups over time by two-way repeated measures ANOVA with Bonferroni correction for pairwise comparisons between individual groups. Depending on data distribution, experimental groups were compared using one-way ANOVA or Kruskal-Wallis, with Posthoc tests (uncorrected Fisher’s LSD or uncorrected Dunn’s test) comparing +/− pectin groups within the normal and high trp feeding groups and comparing normal versus high trp feeding groups within pectin and no pectin feeding groups.

### 16S rRNA gene amplicon sequencing

#### Animal study

DNA was extracted from approximately 250 g cecal and 100 g colonic content using the DNeasy PowerLyzer PowerSoil kit (Qiagen, 12855-100), as described previously^3^, using two blank DNA extraction controls. The V3 region of the 16S rRNA gene was PCR-amplified using 0.2 µl Phusion High-Fidelity DNA polymerase (ThermoFisher Scientific, F-553L), 4 µl HF-buffer, 0.4 µl dNTP (10 mM of each base), 1 µM forward primer (PBU; 5’-A-adapter-TCAG-barcode-CCTACGGGAGGCAGCAG-3’) and 1 µM reverse primer (PBR; 5’-trP1-adapter-ATTACCGCGGCTGCTGG-3’) and extracted DNA diluted to 5 ng/µl in a 20 µl total reaction volume, with a PCR program consisting of initial denaturation for 30s at 98 °C, followed by 30 cycles of 98 °C for 15 s and 72°C for 30 s, and lastly 72 °C for 5 min to allow final extension before cooling to 4 °C. A total of two no-template controls as well as the two DNA extraction controls were included. The PCR products were purified using the HighPrepTM PCR Magnetic Beads (MAGBIO®, AC-60005) with a 96-well magnet stand (MAGBIO®, MyMag 96), according to the manufacturers recommendations. DNA quantity was measured using Qubit® dsDNA HS assay (InvitrogenTM, Q32851) and samples were pooled to obtain equimolar libraries and sequenced on the Ion S5™ System (ThermoFisher Scientific) using Ion OneTouch 2 with the 520 chip kit-OT2 (ThermoFisher Scientific, A27751). Raw sequence reads were analysed as described previously^3^, using CLC Genomic Workbench v8.5 software (CLCbio, Qiagen, Aarhus, DK) to trim off barcodes and primers and the DADA2 pipeline v1.23^49^, according to the tutorial, with few modifications recommended for IonTorrent reads. In brief, reads were quality filtered (maxEE=2, maxN=0, truncQ=2), denoised using pooled data and increased homopolymer gap penalty and band size (pool=TRUE, HOMOPOLYMER_GAP_PENALTY=-1, BAND_SIZE=32) and chimeric sequences were removed and taxonomy was assigned to the resulting ASVs using the RDP 16S rRNA database (v18)^50^. ASVs with less than 100 read counts were removed and relative abundances were calculated by total sum scaling. The top six ASVs represented on average 99.6% (range 99.3-99.8%) of all reads in the colon and cecum samples, and were assigned to Bacteroides (ASV_2, ASV_4, ASV_5, ASV_6), Escherichia (ASV_1) and *Clostridium sensu stricto* (ASV_3). BLAST of the ASV sequences against the 16S rRNA database at NCBI, confirmed 100% identity of the three most abundant ASVs towards *B. thetaiotaomicron* (ASV_2), *E. coli* (ASV_1) and *C. sporogenes* (ASV_3), respectively. BLAST against the 16S rRNA database confirmed Bacteroides classification of ASV_4, ASV_5 and ASV_6, but no 100% match to any species was obtained.

Therefore, these ASVs were additionally searched against the nucleotide collection (GenBank+EMBL+DDBJ+PDB+RefSeq sequences) and was found to match *B. thetaiotaomicron* strain sequences with 100% (ASV_4), 100% (ASV_5) and 98.0% (ASV_6) identity, and these ASVs were collapsed together with ASV_2.

The reaming reads represented either very low abundant ASVs (average relative abundance = 0.39%) matching the same three genus level taxa as the top six ASVs or sporadically detected ASVs (max relative abundance = 0.01%) with high relative abundance in the negative controls (Sum of Sphingomonadaceae, Bradyrhizobium, Rhodopseudomonas, Brevundimonas, Ralstonia, Cutibacterium and Methylobacterium on average 86.9%). The relative abundance of ASV_1 thus represented *E. coli*, the combined relative abundance of ASV_2, ASV_4, ASV_5 and ASV_6 represented *B. thetaiotaomicron* and the relative abundance of ASV_3 represented *C. sporogenes*.

#### Quantitative PCR for total bacterial load

As previously described^3^, we quantified the total bacterial load in cecum and colon samples by quantitative PCR (qPCR) on DNA extracted from these, using universal primers (341F: 5’-CCTACGGGAGGCAGCAG-3’, 518R: 5’-ATTACCGCGGCTGCTGG-3’, final concentration 0.5 µM each) targeting the V3 region of the 16S rRNA gene. Each reaction was performed in triplicates with 2 µl template DNA, the specified primer concentrations and 2X SYBR Green I Master Mix solution (LightCycler® 480 SYBR Green I Master, Roche). Standard curves were generated from known concentrations of 10-fold serial diluted DNA from B. longum subsp. infantis DSM 20088. Plates were run on the LightCycler® 480 Instrument II with 5 min pre-incubation at 95°C, 45 cycles with 15 sec at 95°C, 15 sec annealing at 60°C and 15 sec at 72°C. Data were analyzed with the LightCycler® 480 Software (v1.5) (Roche).

#### 16S rRNA gene amplicon sequencing for in vitro experiments

DNA extraction and PCR of collected pellet samples from fecal cultures and defined community experiments were done as described above. 16S rRNA gene amplicon data was processed using our in-house pipeline. In brief, raw amplicon sequences were demultiplexed using cutadapt (v. 4.1)^51^, denoised using DADA2 (v. 1.22)^49^ and ASVs classified against rdp_train_set_18^52^. Further processing were done using Phyloseq (v.1.42.0)^53^ in R (v. 4.2) (R Core Team 2022).

### Relative gene expression analysis by RT-qPCR

#### RNA extraction

1 ml of bacterial culture was harvested, immediately mixed with two volumes of RNAProtect Bacteria (Qiagen) and pelleted according to the manufacturer’s instructions. The stabilized cell pellets were stored at −80°C until RNA extraction. RNA was extracted using a combination of enzymatic lysis, bead-beating in hot TRIzol, and on-column purification. Briefly, the stabilized pellets were enzymatically lysed for 30 min in a lysozyme solution (15 mg/ml in TE buffer; L4919 Sigma-Aldrich, Merck) combined with 1:10 (v/v) proteinase K (Qiagen) treatment. Pellets from early exponential cultures were lysed in a total volume of 220 µl while the 24h samples were lysed in a total volume of 660µl split into three separate aliquots of 220µl each due to the increased sample material. The lysed cells, were then mixed with 1 ml of TRIzol reagent (Invitrogen) and ∼50mg of glass beads (Ø 0.1 mm, Qiagen), incubated at 65°C for 5 min and beaten for 5 min in a bead beater (Qiagen) set at high speed. After the beating, 200 µl of chloroform was added and the samples were shaken vigorously to mix the phases. Proper phase separation was ensured by centrifugation of the samples at 18,000g for 15’ at 4°C. 700 µl of the resulting RNA containing aqueous phase was subsequently transferred to a new tube, mixed with 500 µl of ethanol (80% v/v), and spin column purified using an RNeasy mini kit (Qiagen) according to the manufacturer’s instructions. The three aliquots originating from the same 24h sample were loaded on the same column. During column purification, on-column DNase I (Qiagen) treatment was included as suggested by the kit manufacturer to remove any trace of genomic DNA. RNA was finally eluted in 50 µl of nuclease-free water. Concentration of the eluted RNA was measured using the QUBIT RNA broad range assay (Invitrogen), purity (A260/A280 and A260/A230 ratios) was estimated using a Nanodrop spectrophotometer (Thermo Fisher Scientific) and integrity was investigated by visual inspection using agarose gel electrophoresis (E-gel EX 1%, Invitrogen). All RNA samples passed the quality control and was stored at −80°C.

#### cDNA synthesis

cDNA was synthesized from 1000 ng of RNA using the GoScript Reverse Transcriptase kit (Promega) according to the manufacturer’s description with random hexamer primers and a final MgCl2 concentration of 5 mM. Identical reactions without the reverse transcriptase were included as negative controls for the qPCR. The cDNA was diluted 10 times with nuclease-free water and stored at −20°C until use.

#### qPCR primer design

The primers used for gene expression analysis are listed in supplementary table 2. Nucleotide sequences of the target genes were retrieved from the genome sequences of the organisms. Primers were designed using software from Integrated DNA Technologies. PrimerQuest with standard settings was used to identify potential amplicons and corresponding primer pairs. The primer pairs were then analysed for possible hairpin formation and primer dimer formation using Oligo Analyzer. Primer specificity was ensured using NCBI Primer Blast^54^ running primer sequences against a custom database comprised of the Genbank entries for the genomes of the strains employed in the defined community experiments. qPCR test runs (see next paragraph) were conducted to ensure that all primer pairs displayed an amplification efficiency above 80% and were free of primer dimer formation and spurious off-target amplification as judged from melting curve analysis.

#### qPCR

qPCR was performed using the intercalating dye based GoTaq qPCR master mix kit (Promega). Briefly, cDNA from an initial RNA input of 10 ng was analysed in a total sample volume of 12 µl with primer concentrations of 800 nM. Samples were mixed in a 384-well PCR plate in technical triplicates. A single replicate of the no-reverse-transcriptase controls as well as a single replicate of a no-template control were included for all samples and amplicons. Assays were run on a Roche LightCycler 480 qPCR machine using a 40 cycle standard two-step PCR protocol with a combined annealing and amplification step at 60°C for 1 min. The qPCR protocol was completed by generation of a melting curve.

#### Data analysis and statistics

Melting curve analysis was performed for all assays after their completion to ensure amplification specificity. The raw fluorescence data was analysed using LinRegPCR^55^. This provided a starting concentration of the amplicon (N0) of each qPCR sample (expressed in arbitrary fluorescence units) calculated from the mean amplification efficiency of each amplicon across all samples, the calculated fluorescence threshold, and the corresponding quantification cycle^56^. The N0 values were used as the basis for the relative expression analysis. *dnaG*, *gyrA*, and *secA* were included as reference genes for all three members of the defined community.

NormFinder^57^ analysis was then performed to select the two reference genes for each individual member that showed the most stable expression level across sample groups. The selected reference genes were then used for normalization to obtain expression ratios for each sample and target gene. Data are presented as fold-change of the expression ratios relative to a reference condition. Unpaired two-tailed t-tests were performed on the expression ratios to determine the statistical significance of the relative expression differences. P<0.05 was considered significant.

### Colorimetric indole measurement using Kovac’s reagent

The bacterial cultures were centrifuged at 14000rpm for 10min at 4°C and the supernatant was collected. 250 µl of supernatant was collected in a new 1.5ml tube and 250 µl of Kovac’s reagent (Sigma-Aldrich, Millipore) were added. The samples were vortexed and incubated at room temperature for 10 min, fast spin (approx. 30 seconds) before the top 100-200µl layer was moved to a 96-well plate, and OD_530nm_ was measured. Standards (0, 10, 20, 50 and 100µM) of Indole (Sigma-Aldrich), in triplicates, were prepared in the same culture media as that of culture supernatants and processed similar to the samples to generate calibration curve. Each day analysis were quantified using the standard curve made on the same day. For quality control, we used six tryptophan metabolites and found that only indole reacts with the Kovac’s reagent (Supplementary Fig. 6).

### Metabolite extraction and profiling

#### Chemicals

Authentic standards of the AAAs and derivatives were obtained from Sigma Aldrich, whereas isotope-labelled AAAs used as internal standards (L-Phenylalanine (ring-d5, 98%), L-Tyrosine (ring-d4, 98%), L-Tryptophan (indole-d5, 98%) and indoleacetic acid (2,2-d2, 96%)) of the highest purity grade available were obtained from Cambridge Isotope Laboratories Inc.

#### Extraction of metabolites from in vitro fermentation samples for AAA metabolite profiling

Culture supernatants from *in vitro* fermentations were thawed at 4°C and then centrifuged at 16.000xg at 4°C for 10 minutes. Subsequently, 80 µL was transferred to a new tube and 20 µL internal standard (40 µg/mL) plus 300 µL acetonitrile were added. These samples were vortexed for 10 seconds and left at −20°C for 10 minutes in order to precipitate the proteins. Then, samples were centrifuged at 16.000xg, 4°C for 10 minutes before 50 µL supernatant of each sample was diluted with 50 µL of sterile water and transferred to a liquid chromatography vial (equalling a 1:10 dilution of the sample with internal standards having a concentration of 1 µg/mL).

#### Extraction of metabolites from serum samples for AAA metabolites profiling

Serum metabolites were extracted as described earlier^58^. Briefly, serums were thawed at room temperature. 10 µl of internal standards (4µg/ml) were added into 40 µl of serum. 50 µl of 0.1 % formic acid was added into serum, vortexed and then 400 µl of cold methanol was added and mixed again by vortex. The samples were then incubated at −20 °C for at least 1 hr for protein precipitation. Samples were then centrifuged twice at 16000g at 4 °C for 10 minutes each to obtain a clear extract which is then dried under nitrogen gas at 40 °C. The sample was then reconstituted into pure sterile 40 µl milliq water and centrifuged again at 5000g at 4 °C for 5 minutes to obtain a clear extract and transferred to a liquid chromatography vial for analysis (equalling a no dilution of the sample with internal standards having a concentration of 1 µg/mL).

#### AAA metabolite profiling in vitro samples and in serum

AAAs and catabolites were semi-quantified *in vitro* samples and in serum by ultra-performance liquid chromatography mass spectrometry (UPLC-MS) using isotopic internal standards with similar molecular structures as previously published^3^. In brief, the samples (2 µL of each) were analysed in random order, however with all samples of the same individual analysed on the same day by a quadrupole time-of-flight mass spectrometry (UPLC-QTOF-MS) system consisting of Dionex Ultimate 3000 RS liquid chromatograph (Thermo Scientific) coupled to a Bruker maXis time of flight mass spectrometer equipped with an electrospray interphase (Bruker Daltonics) operating in positive mode. The analytes were separated on a Poroshell 120 SB-C18 column with a dimension of 2.1×100 mm and 2.7 μm particle size (Agilent Technologies) as previously published^3^. Standard mix solutions (0, 0.8 μg/mL, 2 μg/mL and 4 μg/mL) were analysed as described below. A quality control (QC) was done by taking standard mix solutions of all the analytes (2 µg/mL) in the culture medium and processed similar to the culture supernatant samples to normalize against any loss of the analytes during the processing. In addition, QC samples and standard mix solutions were analysed before and after all the samples and after every 10 samples two standards were analysed and data were processed using QuantAnalysis version 2.2 (Bruker Daltonics) and a calibration curve (fitted to a quadratic regression) with all standards analysed for each metabolite. The calibration curves were established by plotting the peak area ratios of all of the analytes with respect to the internal standard against the concentrations of the calibration standards.

## Supporting information

Supplementary Information_Sinha et al. 2023

## Acknowledgements

We are grateful to Prof. Egon Bech Hansen (DTU) for valuable suggestions and discussions during the work. We thank Marlene Danner Dalgaard at the DTU in-house facility (DTU Multi-Assay Core, DMAC) for performing the 16S rRNA gene sequencing and MS-Omics (Hørsholm, Denmark) for performing the targeted quantitative metabolomics study. We thank Chrysoula Dimopoulou to help with the serum metabolites extraction protocol. Finally, we are very grateful to Bodil Madsen and Katja Ann Kristensen for excellent technical support in the laboratory.

