## Supplementary Information_Sinha et al. 2023 for "Substrate availability and dietary fibre regulate metabolism of tryptophan by human gut microbes"

The supplementary file includes

**Supplementary figures**

- Supplementary Figure 1- Gut bacteria *Clostridium sporogenes* and *Peptostreptococcus anaerobius* produce tryptophan metabolites.
- Supplementary Figure 2- Effect of glucose and tryptophan supplementation on production of tryptophan metabolites.
- Supplementary Figure 3- Carbohydrates supplementation inhibits indole production and stimulates ILA production by infant gut microbiota.
- Supplementary Figure 4- Fibre (or simple carbohydrates) supplementation inhibits indole production by repressing *tnaA* gene expression.
- Supplementary Figure 5- Supplementary data of the effect of tryptophan and fibre supplementation on tryptophan metabolites production by a defined community in vivo.
- Supplementary Figure 6- Qualitative analysis of indolic compounds using Kovac's assay

**Supplementary tables**

- Supplementary Table 1-Diet formula for *in vivo* experiments
- Supplementary Table 2-List of RT-qPCR primers

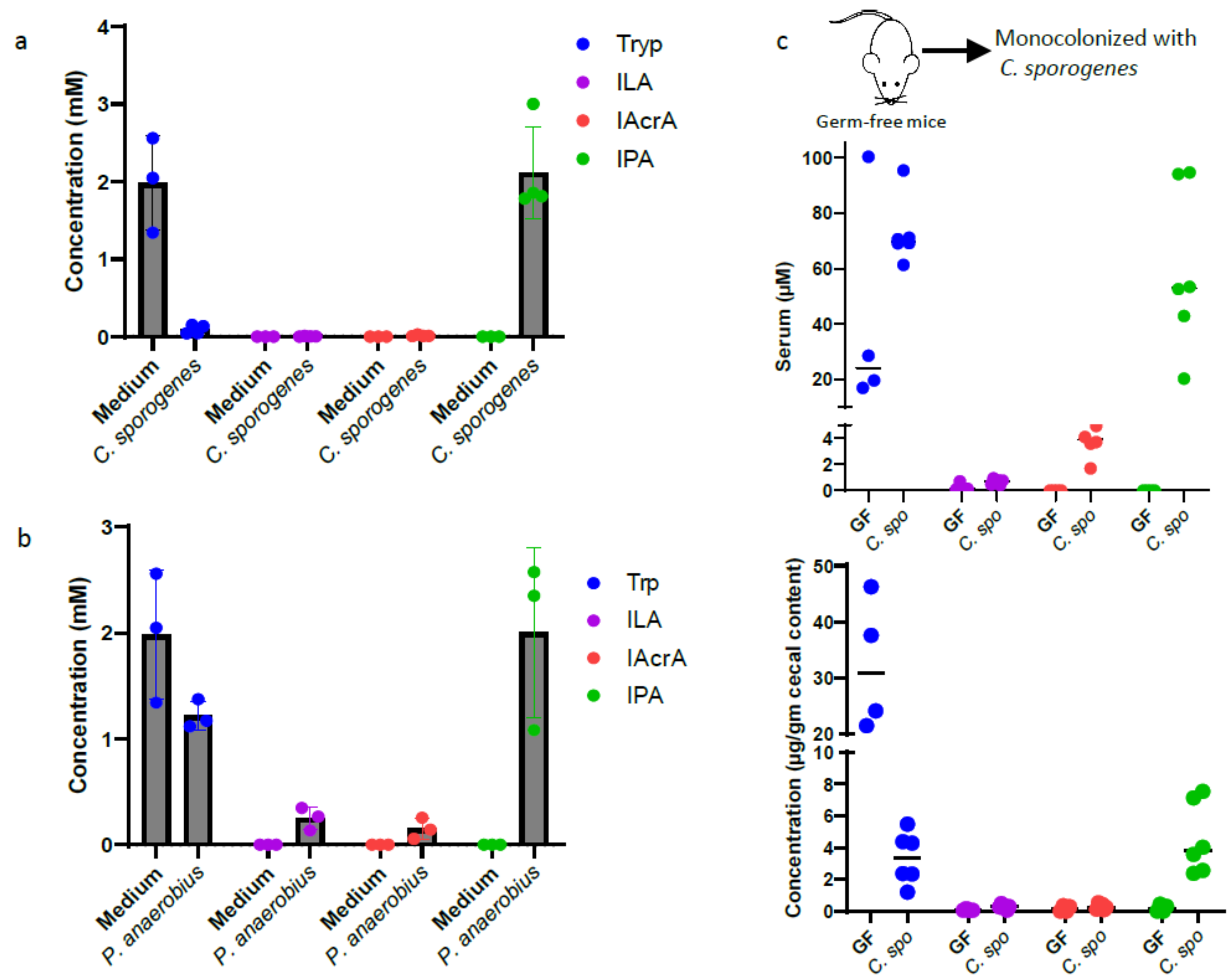


**Supplementary Figure 1. Gut bacteria *Clostridium sporogenes* and *Peptostreptococcus anaerobius* produce tryptophan metabolites.** (a) Tryptophan metabolites in the culture supernatant of *C. sporogenes* grown for 48 hrs in mGAM medium. (b) Tryptophan metabolites in the culture supernatant of *P. anaerobius* grown for 48 hrs in mGAM medium. (c) Tryptophan metabolites in serum and in cecum of germ-free and gnotobiotic mice monocolonized with *C. sporogenes*.

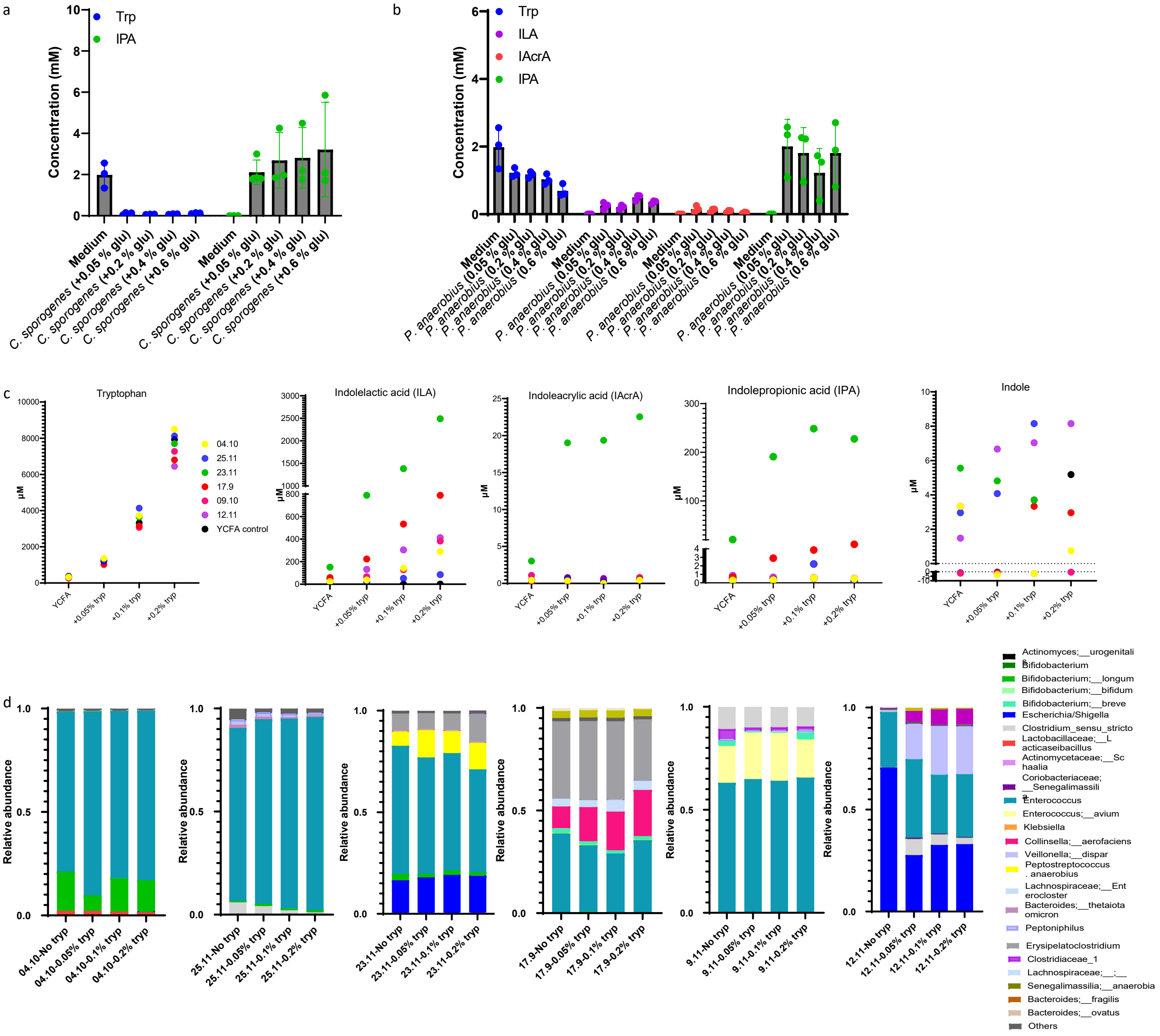


**Supplementary Figure 2. Effect of glucose and tryptophan supplementation on production of tryptophan metabolites.** (a) Tryptophan metabolites accumulation in the culture supernatant of *C. sporogenes* grown in mGAM medium supplemented with final concentration of 0.05, 0.2 or 0.4 and 0.6 % glucose. (b) Tryptophan metabolites accumulation in the culture supernatant of *P. anaerobius* grown in mGAM medium supplemented with final concentration of 0.05, 0.2 or 0.4 and 0.6 % glucose. (c) Absolute concentration of tryptophan metabolites in the culture supernatant of fecal microbiota are shown here. Six infant fecal microbiota are cultured either in YCFA medium or YCFA supplemented with 0.05 or 0.1 or 0.2 % of free tryptophan. (d) 16S rRNA gene sequencing profile of six infant fecal microbiota of the samples.

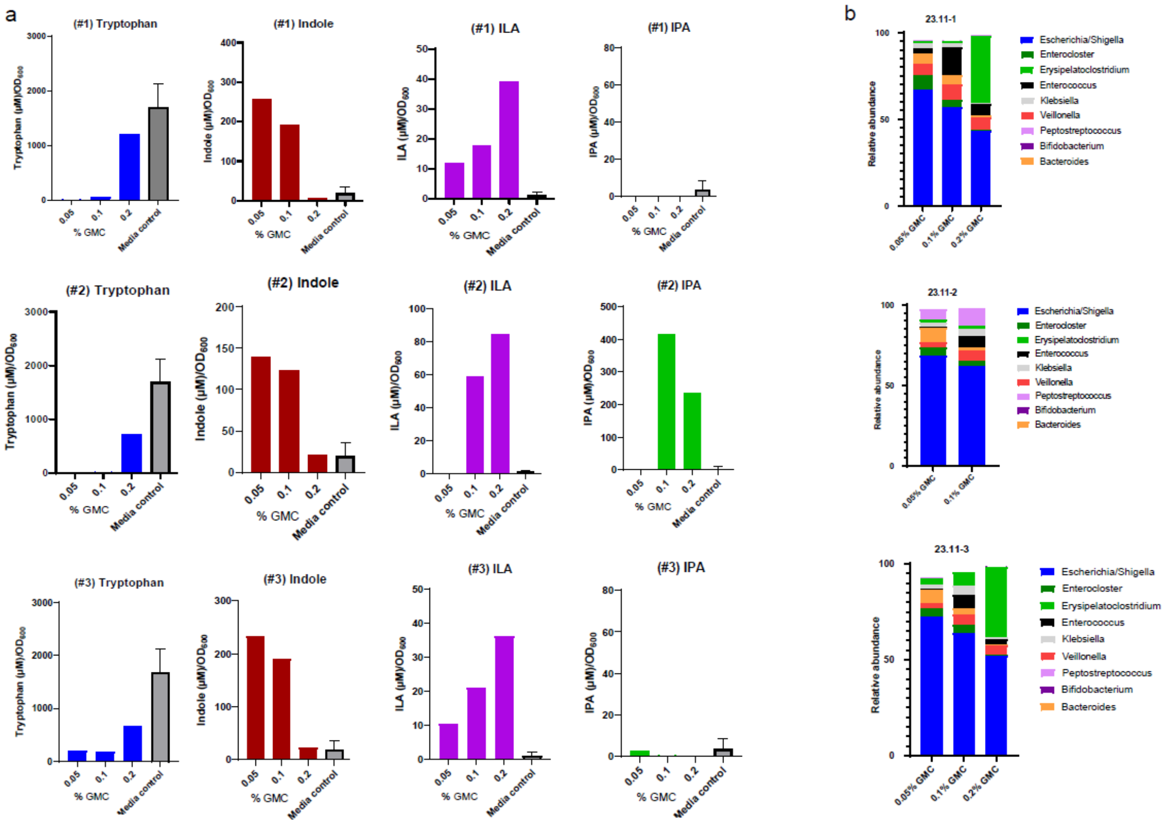


**Supplementary Figure 3. Carbohydrates supplementation inhibits indole production and stimulates ILA production by infant gut microbiota.** (a) Tryptophan metabolites in the culture supernatant of fecal microbiota in YCFA medium supplemented either with 0.05 or 0.1 or 0.2 % each of glucose (G), maltose (M) and cellobiose (C) (collectively referred here as GMC) . One infant fecal sample (23.11) is selected to culture in replicates for analysis since it contained *P. anaerobius*, a bacterium capable of doing Stickland fermentation, and *E. coli*, an indole producer. Metabolites in the individual culture supernatant were normalized against final $OD_{600}$ of the culture. All individual replicates are shown here. (b) 16S rRNA profile of all replicates. Note that Peptostreptpcoccus is frequently lost thus preventing IPA accumulation in many samples.

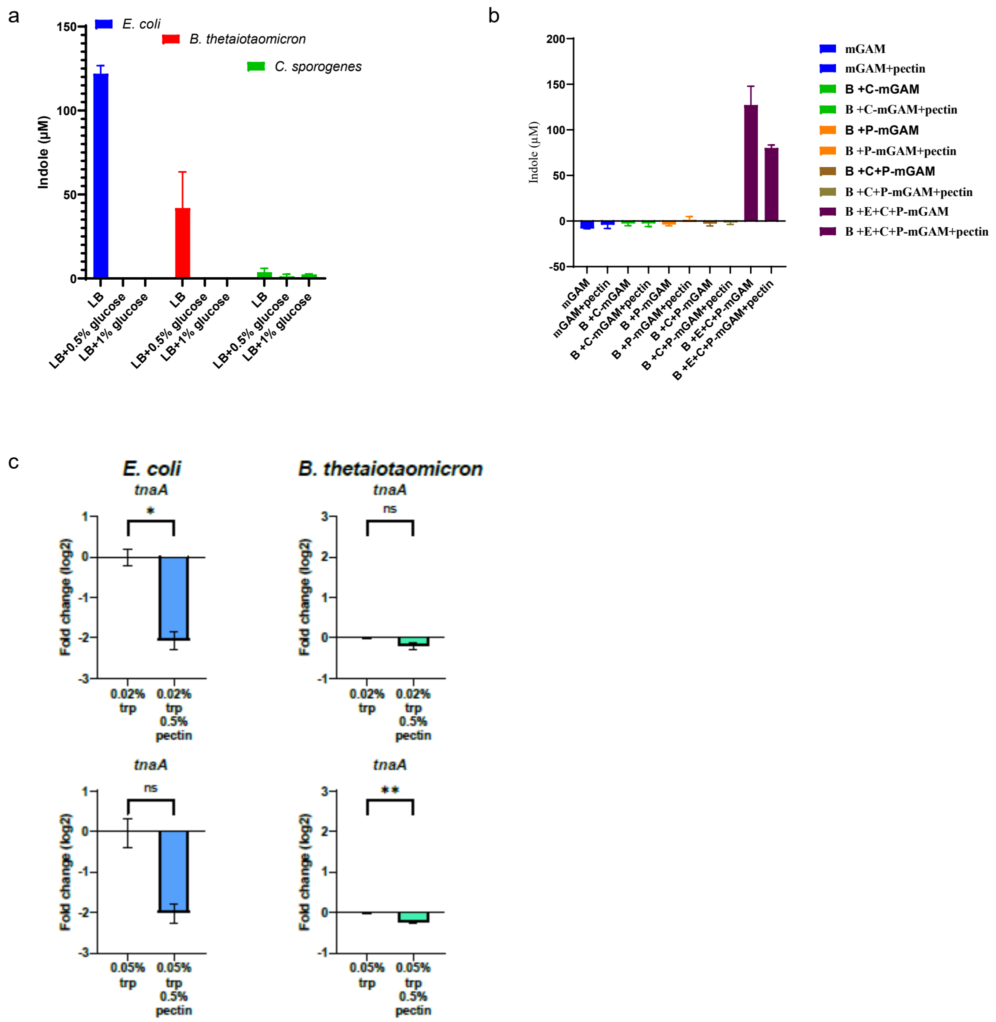


**Supplementary Figure 4. Fibre (or simple carbohydrates) supplementation inhibits indole production by repressing *tnaA* gene expression.** (a) *E. coli*, *B. theta* and *C. spo* are cultured individually either in LB or LB supplemented with 0.5 % glucose or 1 % glucose. Indole production was measured in the culture supernatants. (b) The defined community of combination of *E. coli* (E), *B. theta* (B), *P. anaerobius* (P) and *C. spo* (C) are cultured either in mGAM or in mGAM supplemented with 0.5 % apple pectin. Indole production was measured in the culture supernatants. (c) RT-qPCR to measure *tnaA* mRNA levels in *E. coli* and in *B. theta* in response to pectin supplemented in the growth medium after 24 hrs fermentation.

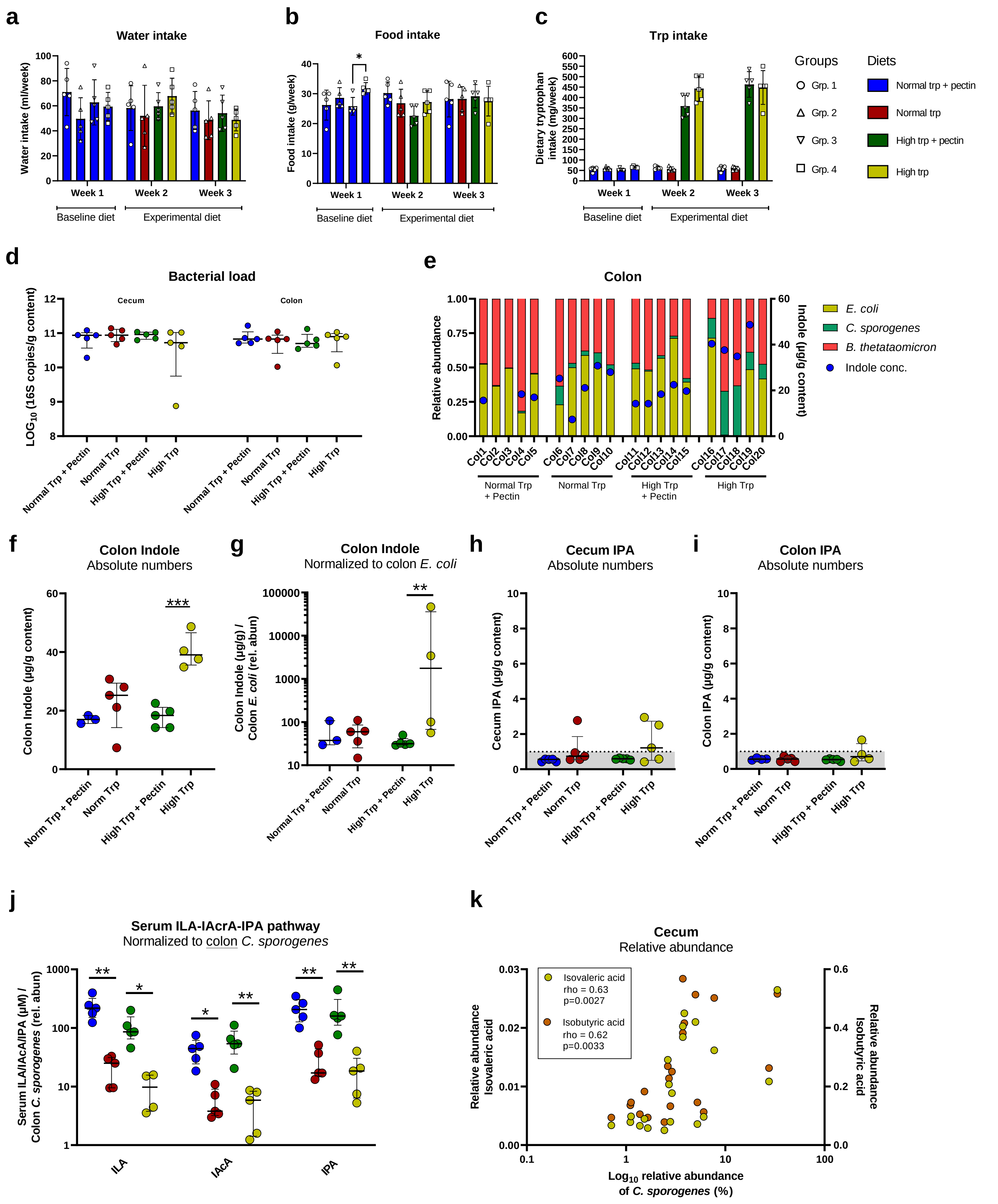
a
Water intake
Water intake (ml/week)
Week 1
Week 2
Week 3
Baseline diet
Experimental diet
b
Food intake
Food intake (g/week)
c
Trp intake
Dietary tryptophan intake (mg/week)
Groups
Grp. 1
Grp. 2
Grp. 3
Grp. 4
Diets
Normal trp + pectin
Normal trp
High trp + pectin
High trp
d
Bacterial load
Cecum
Colon
LOG10 (16S copies/g content)
Normal Trp + Pectin
Normal Trp
High Trp + Pectin
High Trp
e
Colon
Relative abundance
Indole (µg/g content)
E. coli
C. sporogenes
B. thetataomicron
Indole conc.
Normal Trp + Pectin
Normal Trp
High Trp + Pectin
High Trp
f
Colon Indole
Absolute numbers
Colon Indole (µg/g content)
Norm Trp + Pectin
Norm Trp
High Trp + Pectin
High Trp
g
Colon Indole
Normalized to colon E. coli
Colon Indole (µg/g) / Colon E. coli (rel. abun)
h
Cecum IPA
Absolute numbers
Cecum IPA (µg/g content)
i
Colon IPA
Absolute numbers
Colon IPA (µg/g content)
j
Serum ILA-IAcrA-IPA pathway
Normalized to colon C. sporogenes
Serum ILA/IAcA/IPA (µM) / Colon C. sporogenes (rel. abun)
ILA
IAcA
IPA
k
Cecum
Relative abundance
Isovaleric acid
rho = 0.63
p=0.0027
Isobutyric acid
rho = 0.62
p=0.0033
Relative abundance Isovaleric acid
Relative abundance Isobutyric acid
Log10 relative abundance of C. sporogenes (%)

**Supplementary Figure 5. Supplementary data of the effect of tryptophan and fibre supplementation on tryptophan metabolites production by a defined community *in vivo*.** (a) Weekly water intake, (b) food intake and (c) tryptophan intake from individual mice in the different experimental groups described in figure 5a. Bars and error bars indicate mean +/- s.d. No significant differences were found between groups in water intake ($p_{group\_overall} = 0.61$) or food intake ($p_{group\_overall} = 0.39$) according to Two-way repeated measures ANOVA ($p > 0.05$ for all pairwise comparisons between feeding groups at week 2 and 3 after Bonferroni correction). Tryptophan intake differed significantly between groups ($p_{group\_overall} < 0.0001$) and were significantly higher for group 3 and 4 compared to group 1 and 2 in both week 2 and 3 ($p < 0.01$ for all pairwise comparisons across, but not within, trp feeding groups after Bonferroni correction). (d) Total bacterial load quantified using qPCR, showing median and IQR. No significant differences were found using Kruskal-Wallis tests. (e) 16S rRNA gene sequencing profiles show the composition of the defined community in colon of each individual mouse, with indole values in the colon overlaid. Indole values are missing for few mice due to the extremely low samples were present in their colon. (f) Absolute indole concentration in the colon. (g) Indole concentration in the colon, normalized to the relative abundance of *E. coli* in colon. (h-i) Absolute abundance of IPA in (h) cecum and in (i) colon, with grey shaded area indicating background noise (j) Serum tryptophan metabolites normalized to *C. sporogenes* relative abundance in colon. (k) Scatter plot and Spearman's rank correlation of relative isovaleric acid and isobutyric acid versus *C. sporogenes* relative abundance in cecum. All graphs in panel f-j show median and IQR and statistical analysis was done across groups within each metabolite measured using One-way ANOVA (panel f) or Kruskal Wallis tests (panel g-I, k), using uncorrected Fisher's LSD or Dunn's posthoc tests to compare between individual groups. $^*P < 0.05$, $^{**}P < 0.01$, $^{***}P < 0.001$. For panel j, one value for ILA was excluded as an extreme outlier (Grubbs test, alpha < 0.01).

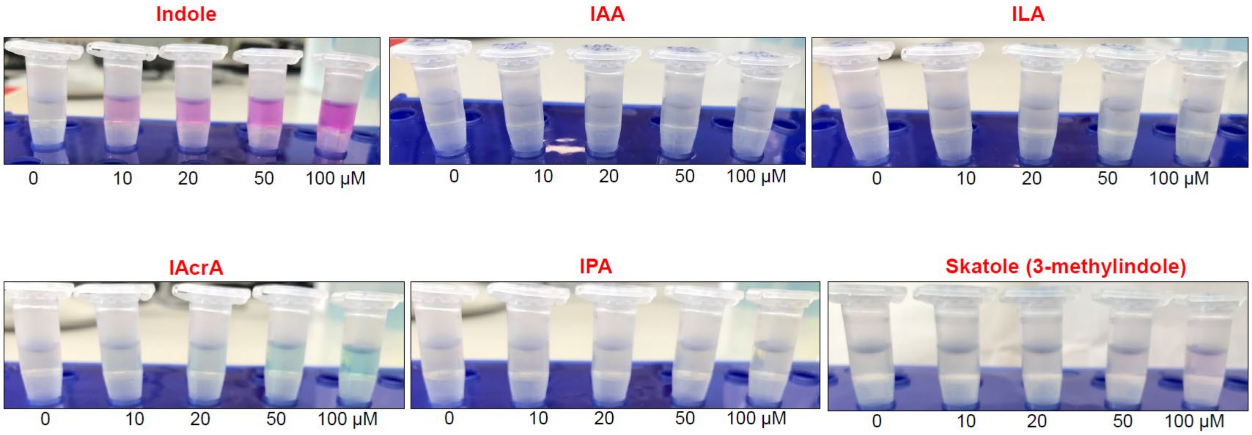


**Supplementary Figure 6. Qualitative analysis of indolic compounds using Kovac's assay** (a) Kovac's reagent specificity against 0, 10, 20, 50 and 100 µM concentration of Indole, IAA, ILA, IAcrA, IPA and skatole were assessed and showed that it is highly specific for indole.

**Supplementary Table 1**-Diet formula for *in vivo* experiments

| Ingredients | Diet 1 A22033102-1.5V | Diet 2 A18041301R-1.5V | Diet 3 A22033103-1.5V | Diet 4 A22033101-1.5V |
|---|---|---|---|---|
| | **(Normal Trp + Pectin)** | **(Normal Trp)** | **(High Trp + Pectin)** | **(High Trp)** |
| Total L-amino acids (g) | 177.1 | 177.1 | 191 | 191 |
| *L-Arginine (g)* | *5.9* | *5.9* | *5.9* | *5.9* |
| *L-Histidine-HCl-$H_2O$ (g)* | *4.5* | *4.5* | *4.5* | *4.5* |
| *L-Isoleucine (g)* | *7.5* | *7.5* | *7.5* | *7.5* |
| *L-Leucine (g)* | *15.7* | *15.7* | *15.7* | *15.7* |
| *L-Lysine-HCl (g)* | *13.1* | *13.1* | *13.1* | *13.1* |
| *L-Methionine (g)* | *5* | *5* | *5* | *5* |
| *L-Phenylalanine (g)* | *8.4* | *8.4* | *8.4* | *8.4* |
| *L-Threonine (g)* | *7.1* | *7.1* | *7.1* | *7.1* |
| ***L-Tryptophan (g)*** | ***2.1*** | ***2.1*** | ***16*** | ***16*** |
| *L-Valine (g)* | *9.2* | *9.2* | *9.2* | *9.2* |
| *L-Alanine (g)* | *5* | *5* | *5* | *5* |
| *L-Asparagine-$H_2O$ (g)* | *7* | *7* | *7* | *7* |
| *L-Aspartic acid (g)* | *5* | *5* | *5* | *5* |
| *L-Cystine (g)* | *4.2* | *4.2* | *4.2* | *4.2* |
| *L-Glutamic acid (g)* | *20.7* | *20.7* | *20.7* | *20.7* |
| *L-Glutamine (g)* | *17.1* | *17.1* | *17.1* | *17.1* |
| *Glycine (g)* | *3* | *3* | *3* | *3* |
| *L-Proline (g)* | *17.6* | *17.6* | *17.6* | *17.6* |
| *L-Serine (g)* | *9.9* | *9.9* | *9.9* | *9.9* |
| *L-Tyrosine (g)* | *9.1* | *9.1* | *9.1* | *9.1* |
| Corn Starch (g) | 372.486 | 397.486 | 358.586 | 383.586 |
| Maltodextrin 10 (g) | 132 | 132 | 132 | 132 |
| Sucrose (g) | 102.0777 | 102.0777 | 102.0777 | 102.0777 |
| Cellulose (g) | 50 | 50 | 50 | 50 |
| **Pectin (g)** | **50** | **0** | **50** | **0** |
| Soybean Oil (g) | 70 | 70 | 70 | 70 |
| t-butylhydroquinone (g) | 0.014 | 0.014 | 0.014 | 0.014 |
| Mineral S10022G (g) | 0 | 0 | 0 | 0 |
| Mineral S10022C (g) | 3.5 | 3.5 | 3.5 | 3.5 |
| Calcium Carbonate (g) | 7.34 | 7.34 | 7.34 | 7.34 |
| Potassium Citrate, 1x$H_20$ (g | 2.4773 | 2.4773 | 2.4773 | 2.4773 |
| Potassium Monobasic (g) | 6.86 | 6.86 | 6.86 | 6.86 |
| CalciumPhosphate,dibasic ( | 7 | 7 | 7 | 7 |
| SodiumChloride (g) | 2.59 | 2.59 | 2.59 | 2.59 |
| SodiumBicarbonate (g) | 7.5 | 7.5 | 7.5 | 7.5 |
| Vitamin Mix V10037 (g) | 15 | 15 | 15 | 15 |
| Choline Bitrartrate (g) | 2.5 | 2.5 | 2.5 | 2.5 |
| Dye Red (g) | 0 | 0.05 | 0 | 0 |
| Dye Yellow (g) | 0 | 0 | 0.025 | 0.05 |
| Dye Blue (g) | 0.05 | 0 | 0.025 | 0 |
| Total (g) | 1008.495 | 983.495 | 1008.495 | 983.495 |

| | **Diet 1** | **Diet 2** | **Diet 3** | **Diet 4** |
|---|---|---|---|---|
| Calories (kcal) | 3925 | 3925 | 3925 | 3925 |
| Tryptophan (g/kg) | 2.08231077 | 2.135242172 | 15.86522491 | 16.26851179 |
| Pectin (g/kg) | 49.57882786 | 0 | 49.57882786 | 0 |
| Tryptophan % | 0.000535032 | 0.000535032 | 0.004076433 | 0.004076433 |
| Pectin % | 0.012738854 | 0 | 0.012738854 | 0 |

**Supplementary Table 2**-List of RT-qPCR primers

| Name | Sequence (5' - 3') | Gene | Reference |
|---|---|---|---|
| *Escherichia coli MG1655 (NCBI Reference Sequence: NC_000913.3)* | | | |
| Ec_tnaA_qPCR-F | GACTGGCTGGCTTATCGTATC | tnaA | This study |
| Ec_tnaA_qPCR-R | GTTTACCGGCATCAACGAATG | | This study |
| Ec_araA_qPCR-F1 | CATCTCGGTATTGGTGGTAAGG | araA | This study |
| Ec_araA_qPCR-R1 | TGTCGATGCAGTTAACCAGTAG | | This study |
| Ec_araF_qPCR-F2 | ATTGGCTGATCGTCGGTATG | araF | This study |
| Ec_araF_qPCR-R2 | TGCTTTAGACAGTTCGCTCAC | | This study |
| Ec_rhaA_qPCR-F2 | GATGAGGTGATCAGCGAGAAG | rhaA | This study |
| Ec_rhaA_qPCR-R2 | ATTGGAGCCAACCGTGTAG | | This study |
| Ec_rhaT_qPCR-F2 | TACGCTGATGACGCCAATTATC | rhaT | This study |
| Ec_rhaT_qPCR-R2 | GAGTTACAATCCCTACGCCAATC | | This study |
| Ec_xylA_qPCR-F1 | GCTACTGGCACACCTTCTG | xylA | This study |
| Ec_xylA_qPCR-R1 | CTCAAATGCGACATCTGCTTTAC | | This study |
| Ec_xylG_qPCR-F1 | AGGTATCGCCATCATTCATCAG | xylG | This study |
| Ec_xylG_qPCR-R1 | CGTAGCGTCATCAGGTCATAATC | | This study |
| Ec_dnaG_qPCR-F | GAAGGCTATATGGACGTGGTG | dnaG | This study |
| Ec_dnaG_qPCR-R | CGCCGTCATAACAGCAAATG | | This study |
| Ec_gyrA_qPCR-F2 | GTGACAAACGTCGTACTGAAATC | gyrA | This study |
| Ec_gyrA_qPCR-R2 | GATACTTAACGTAGCCCTGGTG | | This study |
| Ec_secA_qPCR-F2 | GCTGGTTCTTCCCGTTTCTAC | secA | This study |
| Ec_secA_qPCR-R2 | CCTGGCTTCATACCCAGTTTAC | | This study |
| *Bacteroides thetaiotaomicron DSM 2079 (NCBI Reference Sequence: NC_004663.1)* | | | |
| Bt_tnaA_qPCR-F | GATGCTGGGTGATGAAAGTTATG | tnaA (BT_RS07555) | This study |
| Bt_tnaA_qPCR-R | CGTCCTTGATGGGTAGGAATG | | This study |
| Bt_gyrA_qPCR-F | CGTGGGTGAAGTATTGGGTAAG | gyrA | This study |
| Bt_gyrA_qPCR-R | GGGCTATCACCATCTACAGAAC | | This study |
| Bt_dnaG_qPCR-F | CTTCGCCCGCAAACATAATTC | dnaG | This study |
| Bt_dnaG_qPCR-R | CGTTTGATCGGATCTCTTCCC | | This study |
| Bt_secA_qPCR-F | CGATGAGGTTGACTCGGTATTG | secA | This study |
| Bt_secA_qPCR-R | GAGCTTCTACCAGACGTTCTAC | | This study |
